# Pulses of RhoA Signaling Stimulate Actin Polymerization and Flow in Protrusions to Drive Collective Cell Migration

**DOI:** 10.1101/2023.10.03.560679

**Authors:** Weiyi Qian, Naoya Yamaguchi, Patrycja Lis, Michael Cammer, Holger Knaut

## Abstract

In animals, cells often move as collectives to shape organs, close wounds, or—in the case of disease—metastasize. To accomplish this, cells need to generate force to propel themselves forward. The motility of singly migrating cells is driven largely by an interplay between Rho GTPase signaling and the actin network (Yamada and Sixt, 2019). Whether cells migrating as collectives use the same machinery for motility is unclear. Using the zebrafish posterior lateral line primordium as a model for collective cell migration, we find that active RhoA and myosin II cluster on the basal sides of the primordium cells and are required for primordium motility. Positive and negative feedbacks cause RhoA and myosin II activities to pulse. These pulses of RhoA signaling stimulate actin polymerization at the tip of the protrusions and myosin II-dependent actin flow and protrusion retraction at the base of the protrusions, and deform the basement membrane underneath the migrating primordium. This suggests that RhoA-induced actin flow on the basal sides of the cells constitutes the motor that pulls the primordium forward, a scenario that likely underlies collective migration in other—but not all (Bastock and Strutt, 2007; Lebreton and Casanova, 2013; Matthews et al., 2008)—contexts.

## Introduction

To move, cells must generate forces to propel themselves forward. Classic studies in cell culture show that mesenchymal cells move in a three-step manner; they extend their front through protrusions, attach to their substrate through focal adhesions (FAs), and contract their rear with resolution of the FAs (Parsons et al., 2010; Webb et al., 2002; Yamada and Sixt, 2019). In part, these steps are coordinated by small Rho GTPases. In the simplest model, Rac and Cdc42 are active in the front of the cells and stimulate actin polymerization to form lamellipodia and filopodia (Nobes and Hall, 1995; Ridley et al., 1992), respectively, while RhoA is mostly active in the body and rear of migrating cells and regulates actomyosin contraction and rear retraction (Jaffe and Hall, 2005; Ridley and Hall, 1992; Ridley et al., 2003). *In vivo* studies have confirmed some aspects of this model for locomotion in singly migrating cells (Pocha and Montell, 2014). However, whether the coordinated movement of cells in a migrating tissue *in vivo* also involves such a three-step manner is largely unclear. To address this question, we used the zebrafish posterior lateral line primordium as a model. The primordium is a tissue of about 140 cells that migrates directly under the skin from behind the ear to the tip of the tail (Nogare and Chitnis, 2017). All the cells in the primordium sense the attractant chemokine Cxcl12a (David et al., 2002; Donà et al., 2013; Venkiteswaran et al., 2013), adhere to each other, and contribute to the collective’s forward movement (Colak-Champollion et al., 2019).

## Results and discussion

### Active RhoA and myosin II localize to puncta in the primordium

Because Rac, Cdc42, and RhoA are critical for cell migration, we assessed their localization using Rho GTPase localization sensors. We inspected the localization of active Rac/Cdc42 and active RhoA using superfolder GFP (sfGFP) fusions to the protein binding domain of PAK1 (Benard et al., 1999) and the AHPH domain of Anillin (Piekny and Glotzer, 2008; Priya et al., 2015; Sun et al., 2015; Tse et al., 2012), respectively. These fusion proteins were expressed in the primordium under the control of the *cxcr4b* promoter on a bacterial artificial chromosome (BAC) transgene (*cxcr4b:PAK-PBD-sfGFP* and *cxcr4b:sfGFP-AHPH*). This BAC contains a 69 kb-long genomic DNA fragment that encompasses the *cxcr4b* locus (Venkiteswaran et al., 2013). In contrast to the localization patterns observed in singly migrating cells (Bros et al., 2019; Fritz and Pertz, 2016), we did not detect elevated levels of active Rac/Cdc42 at the front of the cells in the primordium. Instead, active Rac/Cdc42 localizes fairly uniformly to the membranes of the cells (Fig. S1A). In contrast, active RhoA localizes to specific sites in the cells of the primordium. Consistent with its reported role in neuromast formation(Harding and Nechiporuk, 2012; Olson et al., 2023) active RhoA concentrates at the center of the forming neuromasts (Fig. 1A, asterisks). In addition, we find clusters of active RhoA — which we refer to as active RhoA puncta — at the basal sides of the primordium cells and at the membranes of primordium cells that border the skin-muscle junction — which we refer to as the primordium’s outer border (Fig. 1A, Video 1).

**Figure 1.**
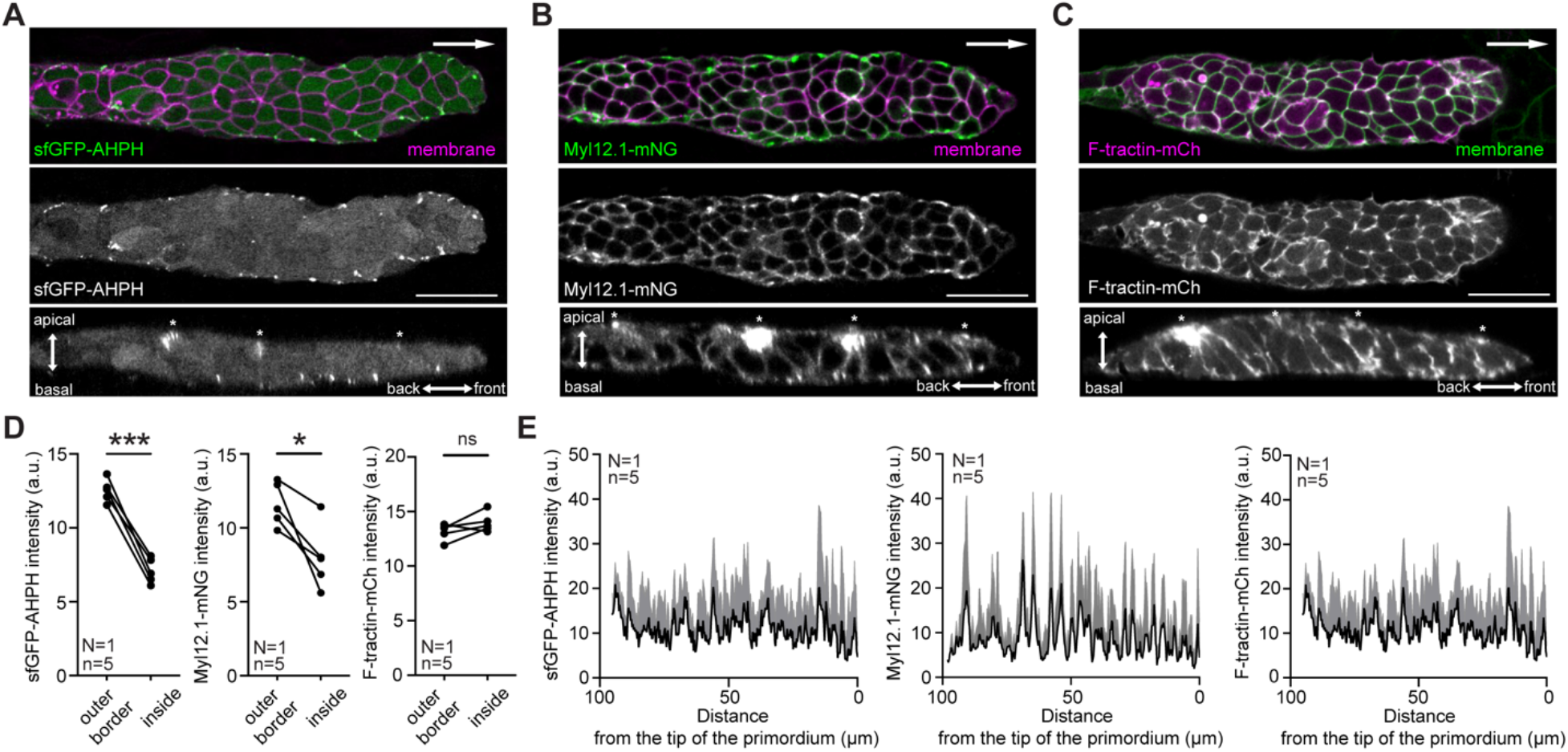
Distribution of RhoA signaling pathway components in the primordium. **A–C**, Images of the localization pattern of sfGFP-AHPH (A), Myl12.1-mNG (B), and F-tractin-mCherry (C) in the primordium. Top and middle panels are single xy-sections and bottom panels are single xz-sections through the middle of the primordium from the z-stacks shown in Video 1. White arrow in top panels indicates direction of primordium migration. Asterisks in lower panels indicate the location of the apical constrictions. Scale bar, 25 µm. **D**, Quantification of mean sfGFP-AHPH (left), Myl12.1-mNG (middle), and F-tractin-mCherry (right) fluorescence intensity at the outer border (0–10 μm margin around the primordium) and inside the primordium (>10 μm from the primordium’s margin). \**P* = 0.022, \*\*\**P* = 0.0003 (Paired t test). **E**, Fluorescence intensity profile of sfGFP-AHPH (left), Myl12.1-mNG (middle), and F-tractin-mCherry (right) along a 10 μm-wide margin of the primordium from the primordium’s front to its back. Black curve represents the mean and grey area represents the mean plus s.d.. In all panels, n indicates number of primordia/embryos, N indicates number of experiments.

The unusual localization pattern of active RhoA on the basal side of the primordium prompted us to investigate the role of RhoA signaling in primordium migration further. RhoA can signal through the Rho kinase (ROCK) to activate non-muscle myosin II and, thus, actomyosin contraction (Clarke and Martin, 2021). We therefore assessed the distribution of non-muscle myosin II (myosin II) and filamentous actin (F-actin) in the primordium using Myl12.1 (Maître et al., 2012) fused to mNeonGreen (mNG) and F-tractin (Schell et al., 2001) fused to mCherry. Both reporter fusion proteins were expressed in the primordium under the control of the *cxcr4b* promoter on a BAC transgene (*cxcr4b:Myl12.1-mNG, cxcr4b:F-tractin-mCherry*). Consistent with the distribution of active RhoA, myosin II and F-tractin are concentrated at the center of the forming neuromasts at the apical side of the primordium (Fig. 1B–C, asterisks). Myosin II also accumulates in puncta on the basal sides of the primordium cells and along the primordium’s outer border (Fig. 1B, Video 1, 2). This localization pattern largely overlaps with the localization pattern of phosphorylated myosin light chain (MLC), indicating that Myl12.1-mNG puncta correspond to clusters of active myosin II (Fig. S1B). In contrast to active RhoA and myosin II, F-actin forms dense networks on the basal sides of the primordium cells (Fig. 1C). Additionally, myosin II and F-actin are enriched at sites of cell-cell contacts (Fig. 1A–C, Movie 1, 2) to which active RhoA does not localize (Fig. 1A). Quantification of the puncta intensities of sfGFP-AHPH and Myl12.1-mNG indicates that RhoA-mediated actomyosin activity is more pronounced along the primordium’s outer border than within the primordium (Fig. 1D) but is fairly evenly distributed along the primordium’s outer border along the front-rear axis (Fig. 1E). However, this pattern is not reflected in the distribution of F-actin which is not preferentially enriched along the primordium’s outer border (Fig. 1D) or the front-back axis (Fig. 1E) probably because many pathways can induce F-actin formation (Bernstein and Bamburg, 2010; Hung et al., 2010; Levayer and Lecuit, 2012; Rottner and Stradal, 2011). While enriched at the primordium’s outer border, active RhoA and myosin II puncta do not form a continuous cable-like structure at the primordium’s outer border as observed in migrating neural crest (Shellard et al., 2018), wound edge cells (Martin and Lewis, 1992) and groups of tumor cells (Hidalgo-Carcedo et al., 2010; Zajac et al., 2018). Thus, in contrast to the uniform distribution of active Rac/Cdc42 along the membranes, active RhoA — together with myosin II — is enriched at the apical constrictions of the forming neuromasts and localizes to puncta at the basal sides and the outer border of the primordium cells.

### The primordium requires RhoA signaling for motility

The localization of active RhoA and myosin II to puncta in the primordium cells suggested that RhoA signaling might be required for primordium motility. To test this idea, we first blocked RhoA signaling using Rockout and blebbistatin, two small molecular inhibitors directed against ROCK and myosin II, respectively. In control-treated embryos, the primordium completed its migration indistinguishable from wild-type embryos at 48 hpf (Fig. S1C–E, Video 3). However, in embryos treated with Rockout or blebbistatin 4 hours after the onset of migration (24 hpf), the primordium slowed down, stalled and on average only completed 50% or 70% of its migration by 48 hpf, respectively (Fig. S1D, E, Video 3). In contrast to primordium cells in blebbistatin-treated embryos, some primordium cells started to die in embryos treated with Rockout for more than 3 hours (Video 3).

Because treatment with Rockout or blebbistatin blocks ROCK or myosin II activity in the entire embryo, we used the LexPR-LexOP system to manipulate RhoA signaling specifically in the migrating primordium with temporal control (Fig. 2A, B). The LexPR-LexOP system consists of the hybrid transcription factor LexPR, the LexOP operator and minimal promoter DNA sequences. Upon binding of the hormone RU-486, LexPR dimerizes and binds to the LexOP operator and induces the transcription of the gene of interest placed under the control of the LexOP operator (Fig. 2A) (Emelyanov and Parinov, 2008; Kenyon et al., 2018). Using a BAC transgene, we expressed LexPR under the control of the *cxcr4b* promoter in the primordium. On the same BAC transgene, 15 kb downstream of the *cxcr4b* locus, we placed the C3 exoenzyme (C3) or constitutively active myosin phosphatase 1 (caMYPT1) under the control of the LexOP operator (*cxcr4b:LexPR-LexOP:C3*, *cxcr4b:LexPR-LexOP:caMYPT1*). C3 ADP-ribosylates and inactivates Rho-like proteins (Boquet, 1999) and caMYPT1 — in complex with the phosphatase PP1 — dephosphorylates and inactivates myosin II (Smutny et al., 2017; Terrak et al., 2004). RU-486 induced expression of C3 in the primordium blocked activation of RhoA in the primordium (Fig. 2C, D), and RU-486 induced expression of caMYPT1 in the primordium blocked myosin II clustering in the primordium (Fig. S2A). This indicates that these transgenic lines can be used to interfere with RhoA signaling in the primordium with temporal control. When we blocked RhoA activation through C3 induction in the primordium, the primordium slowed down and ceased to migrate (Fig. 2E, F, Video 4). Also, cells that started to divide failed to complete division, shed from the collective, and died in primordia expressing C3 (Video 4). In RU-486-treated embryos that did not carry the *cxcr4b:LexPR-LexOP:C3* transgene or that carried the same transgene but with an in-frame deletion in the C3 coding sequence introduced by genome-editing, primordium migration was not affected (Fig. 2E, F, Fig. S1F). A likely reason for cell shedding and cell death in primordia of Rockout-treated embryos and C3-expressing primordia is the requirement of RhoA signaling in constricting the contractile ring during cytokinesis (Basant and Glotzer, 2018). Similar to inducing C3 expression in the primordium, inducing caMYPT1 expression in the primordium also stalled primordium migration but did not lead to cell shedding or cell death (Fig. S1G, Fig. S2E, F, Video 4). From these observations we conclude that RhoA signaling through myosin II is required in the primordium for its motility.

**Figure 2.**
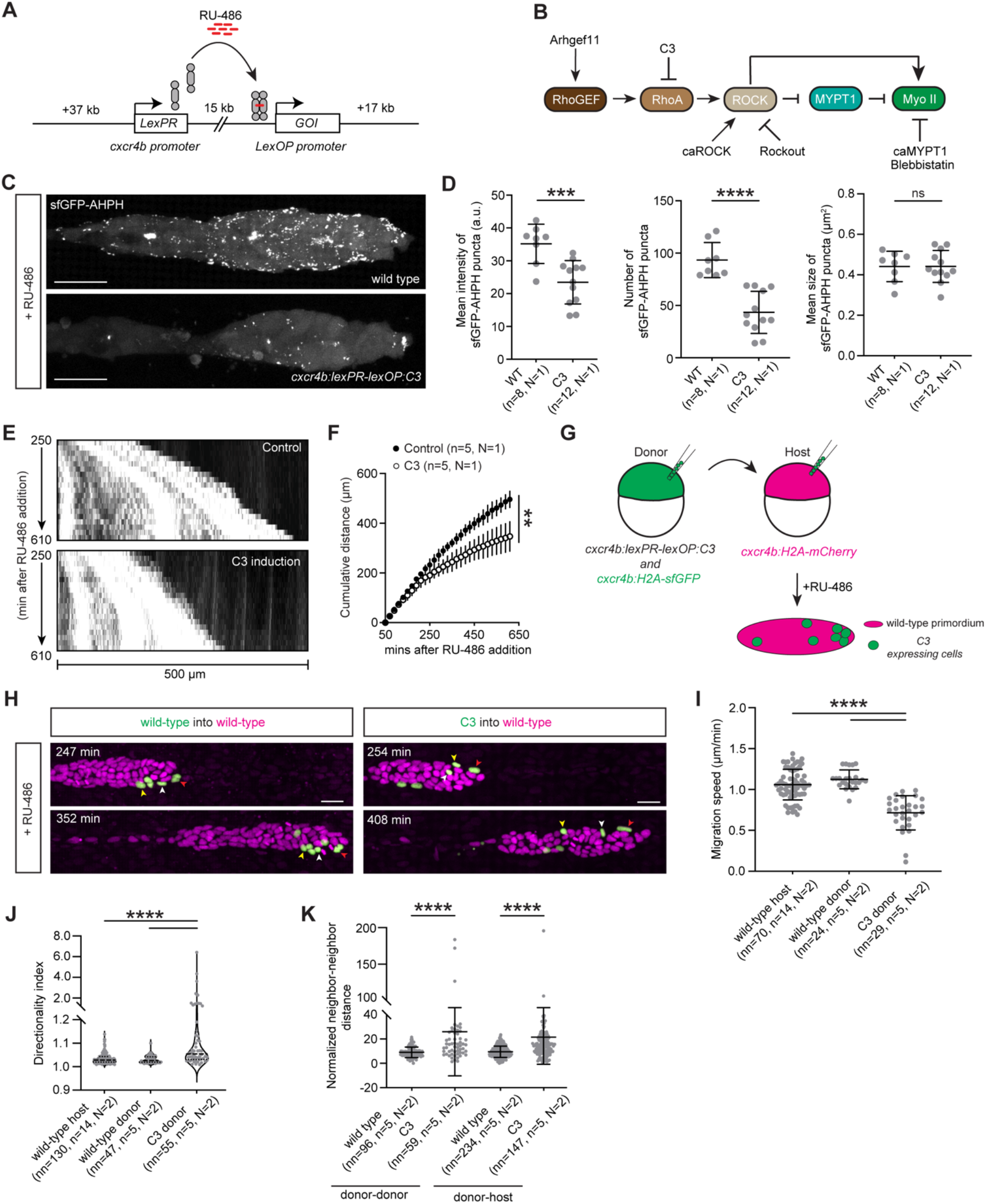
RhoA signaling is required for motility on a tissue and single cell level in the primordium. **A**, Schematic illustration of the *LexPR-LexOP* drug-inducible system expressed under the control of the *cxcr4b* promoter from a 69 kb genomic DNA fragment on a bacterical artificial chromosome. **B**, Schematic illustration of the RhoA/ROCK signaling pathway with inhibitors and regulators used in this study. **C,** Representative images of the sfGFP-AHPH localization pattern in primordia of RU-486-treated control embryo (top) and embryo expressing C3 under the control of the LexOP promoter upon induction by RU-486 (bottom). Images are maximum-projected z-stacks. Scale bar, 25 µm. **D,** Plot of the mean fluorescence intensity, number, and mean size of sfGFP-AHPH puncta shown in C. Mean and s.d. are indicated. \*\*\*\**P* < 0.0001, \*\*\**P =* 0.0008, ns = not significant (Unpaired t test). n indicates number of primordia/embryos, N indicates number of experiments. **E**, Kymographs of time lapse videos of primordia in a control embryo (top) and an embryo expressing C3 in the primordium (bottom). The kymographs trace the tips of the primordia in Video 4. **F**, Plot of the cumulative primordium migration distance in control embryos (black circles) and embryos expressing C3 in the primordium (white circles). Mean and s.d. are indicated. \*\**P* = 0.0012 (Unpaired t test). n indicates number of primordia/embryos, N indicates number of experiments. **G,** Experimental design for the generation of chimeric primordia consisting of mostly wild-type cells and a few cells that are either wild-type or expressing C3. **H**, Representative time lapse images of chimeric primordium consisting of mostly wild-type cell (magenta) and a few cells that are either wild-type (left) or expressing C3 (right, green) shown in Video 5. Arrowheads trace the positions of 3 cells in the primordium over time. Time in min after addition of RU-486. Scale bar, 20 µm. **I**, Quantification of the migration speed for wild-type host cells and donor cells that are either wild-type or expressing C3. Mean and s.d. are indicated. nn indicates number of cells, n indicates number of primordia/embryos, N indicates number of experiments. \*\*\*\**P* < 0.0001 (Kruskal-Wallis test). **J**, Quantification of the directionality index of wild-type host cells, wild-type donor cells, and donor cells expressing C3 in chimeric primordia. Dotted lines indicate the quartiles and dashed lines indicate the medians. nn indicates number of cell tracks, n indicates number of primordia/embryos, N indicates number of experiments. \*\*\*\**P* < 0.0001 (Kruskal-Wallis test). **K**, Quantification of the normalized neighbor-neighbor distances between donor-donor and donor-host cells in chimeric primordia of indicated genotype. The neighbor-neighbor distances were normalized to the average speed of the primordium cells. Mean and s.d. are indicated. nn indicates number of cell pairs, n indicates number of primordia/embryos, N indicates number of experiments. \*\*\*\**P* < 0.0001 (Kruskal-Wallis test).

To test whether over-activation of RhoA signaling in the primordium also perturbs its migration, we generated two transgenic lines that express Arhgef11 (also known as PDZ-RhoGEF) or a constitutively active version of ROCK (caROCK) in the primordium upon RU-486 addition using the LexPR-LexOP system under the control of the *cxcr4b* promotor (*cxcr4b:LexPR-LexOP:Arhgef11, cxcr4b:LexPR-LexOP:caROCK*). Arhgef11 is a Rho guanine nucleotide exchange factor (RhoGEF) that activates RhoA (Müller et al., 2020) and caROCK is truncated version of ROCK that lacks its autoinhibitory domain and activates myosin II (Amano et al., 1999). In Arhgef11-expressing primordia, we observed an increase in the number of active RhoA puncta (Fig. S2C) and the size of myosin II puncta (Fig. S2D) and the primordium ceased to migrate (Fig. S1I, Fig. S2E, F, Video 4). Similarly, in caROCK-expressing primordia the intensity and size of myosin II puncta increased (Fig. S2B) and the primordium migrated more slowly than in control embryos (Fig. S1H, Fig. S2E, F, Video 4). The onset of perturbed myosin II clustering and primordium migration was delayed in caROCK-expressing primordia compared to C3-, caMYPT1-, or Arhgef11-expressing primordia because induction of caROCK expression was delayed; the onset of expression from the LexOP operator in these lines was assessed by the expression of the nuclear-targeted blue fluorescent protein co-expressed using a T2A sequence. Together, these observations suggest that the primordium requires proper RhoA signaling through myosin II for its motility.

### RhoA is required in individual cells for co-migration with the primordium

Our observation that changing RhoA signaling in the primordium impairs its migration suggests that each individual cell in the primordium requires RhoA signaling to migrate. To test this idea, we placed wild-type control or *cxcr4b:LexPR-LexOP:C3* transgenic cells into wild-type primordia by blastomere transplantation, added RU-486, and assessed the migratory behavior of the cells in the primordium. The nuclei of the transplanted donor cells were labeled with GFP and the nuclei of the host primordium cells were labeled with mCherry for cell tracking (Fig. 2G). Using this approach, we found that C3-expressing cells migrate slower and less directional than wild-type donor control or wild-type host neighboring cells (Fig. 2H–J), increase the distance to their wild-type neighbors (Fig. 2K), and fall behind (Video 5). This suggests that RhoA is required in individual primordium cells to co-migrate with wild-type neighboring cells. A similar cell autonomous requirement was reported for primordium cells lacking the guidance receptors Cxcr4 (Colak-Champollion et al., 2019; Haas and Gilmour, 2006). This indicates that wild-type primordium cells are unable to efficiently pull along cells that have impaired motility or directionality, probably because the adhesion between the primordium cells is too weak for wild-type cells to drag such cells along over long distances.

### Pulses of RhoA activation contract the actomyosin network in the primordium

In many contexts, RhoA triggers pulsed actomyosin network contractions that drive tissue morphogenesis (Levayer and Lecuit, 2012). To test whether RhoA signaling is also pulsatile in the primordium, we characterized the dynamics of active RhoA, myosin II, and F-actin accumulation in the primordium cells. For this we generated embryos which expressed either sfGFP-AHPH and Myl12.1-mScarlet (*cxcr4b:sfGFP-AHPH*; *cxcr4b:Myl12.1-mScarlet*), sfGFP-AHPH and F-tractin-mCherry (*cxcr4b:sfGFP-AHPH*; *cxcr4b:F-tractin-mCherry*), or Myl12.1-mNG and F-tractin-mCherry (*cxcr4b:Myl12.1-mNG*; *cxcr4b:F-tractin-mCherry*) in the primordium. These reporters label active RhoA and myosin II, active RhoA and F-tractin, and myosin II and F-actin with green and red fluorescent proteins, respectively (Fig. 3A, Video 6). We imaged the primordia in these double transgenic embryos every 12 seconds for 40 minutes and plotted the fluorescence intensities of the reporters for active RhoA, myosin II, and F-actin along the outer border of the migrating primordium over time (Fig. S3A). The resultant space-time plots — or kymographs — suggested that active RhoA clustering, myosin II clustering, and F-actin accumulation are pulsatile and temporally correlated (Fig. 3B–D). To quantify the pulse period and the temporal correlation, we extracted the fluorescence intensity profiles of individual sfGFP-AHPH or Myl12.1-mNG puncta together with the fluorescence intensity profiles of Myl12.1-mScarlet or F-tractin-mCherry from the kymographs over time, calculated the auto- and cross-correlations, and determined the pulse periods (Fig. S3A). This analysis showed that the average pulse periods for active RhoA, myosin II, and F-actin clusters of 9 ± 4, 8 ± 3 and 8 ± 3 minutes, respectively, are similar to each other (Fig. 3E), are comparable between the front and rear halves of the primordium (Fig. S3D), and are distributed similarly (Fig. S3E–S3G). Also, paired comparison of the pulse periods for any of the reporter combinations revealed no significant differences (Fig. 3F-H). These average pulse periods are longer than the pulse periods of 2 to 7 minutes seen in other contexts (Blanchard et al., 2010; David et al., 2010; He et al., 2010; Martin et al., 2009; Munjal et al., 2015), likely reflecting the slower dynamics of primordium migration. Intriguingly, expressing Arhgef11 in the primordium decreased the pulse period of active RhoA and myosin II cluster to 4 ± 1 minutes (Fig. 3E, S3C). This indicates that RhoGEF levels are limiting the pulse frequency in the primordium, possibly by limiting the rate at which active RhoA generates its own inhibitor, and suggests that RhoGEF levels are a means to tune the frequency of RhoA pulsing, although this may not be the case in all contexts. Consistent with the similar pulse periods of active RhoA, myosin II, and F-actin, the normalized cross-correlation coefficients for any combination of the reporters for active RhoA, myosin II, and F-actin were above 0.8 (Fig. 3I). As positive and negative controls, we determined the cross-correlation coefficients Myl12.1-mNG and Myl12.1-mScarlet — two signals that should be perfectly correlated — and Myl12.1-mNG and membrane-tethered mCherry (Fig. S3B) — two signals that should not be correlated — in the primordium to be 0.9 and 0.2, respectively (Fig. 3I). Together these observations are consistent with the idea that active RhoA, myosin II clusters, and F-actin accumulation oscillate together in the primordium.

**Figure 3.**
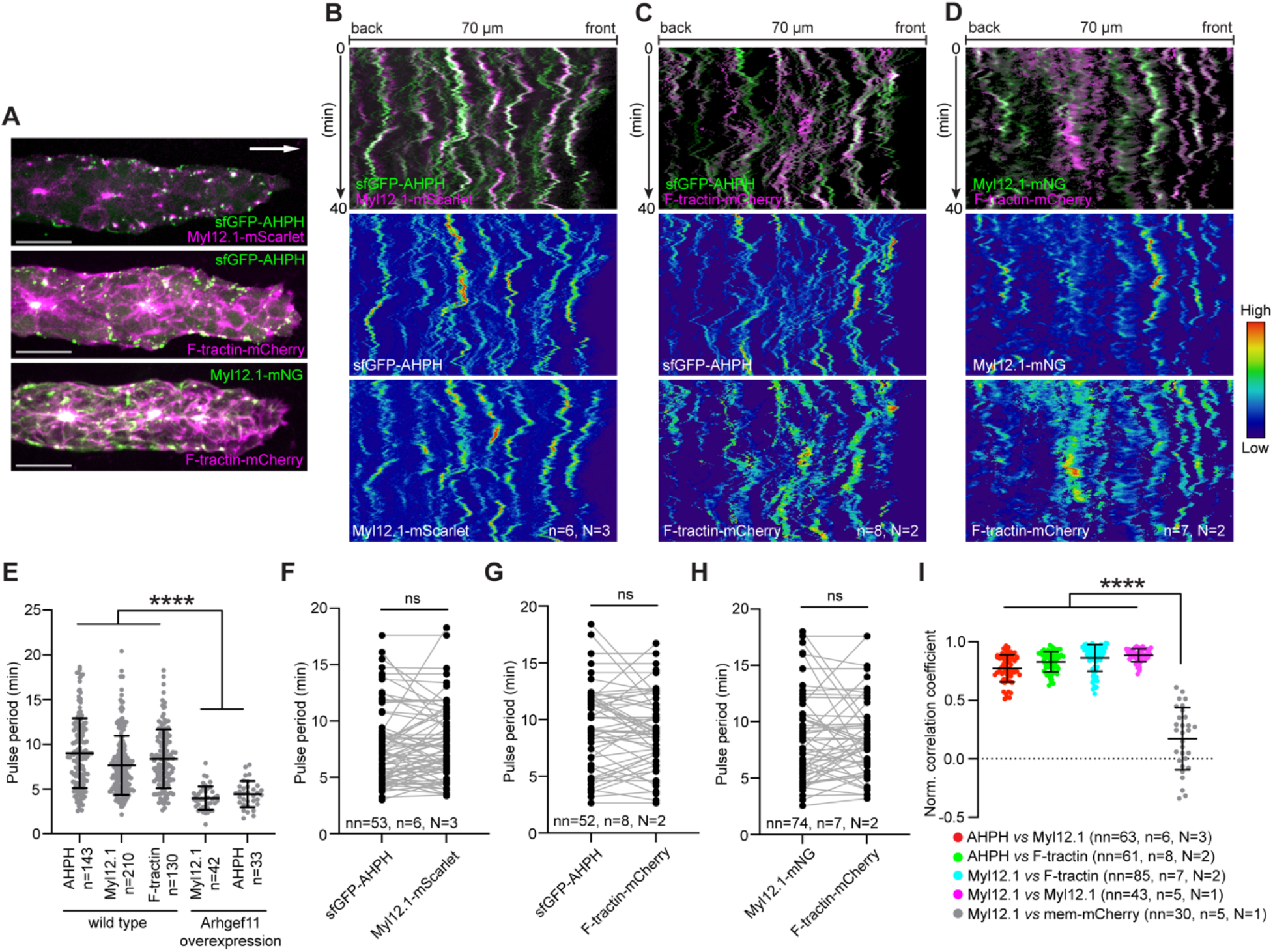
RhoA signaling is pulsatile in the primordium. **A**, Representative images of the localization patterns of AHPH, Myl12.1, and F-tractin reporters in wild-type primordia. Images are maximum-projected z-stacks from a time lapse video (Video 6). Scale bar, 20 µm. Arrow indicates direction of migration. **B–D**, Kymographs along the front-back margin of primordia expressing indicated combinations of fluorescent protein-tagged AHPH, Myl12.1, and F-tractin reporters from 40 min time lapse videos with a temporal resolution of 12 s as detailed in Fig. S3 (Video 6). n indicates number of primordia/embryos, N indicates number of experiments. **E**, Plot of the puncta pulse periods of indicated reporters in wild-type primordia and primordia over-expressing Arhgef11. \*\*\*\**P* < 0.0001 (Kruskal-Wallis test). n indicates number of puncta. For wild-type, primordia number > 14 from 2 or more experiments; for Arhgef11 overexpression, primordia number = 5 from 1 experiment. **F–H**, Plot of the pulse periods of puncta of indicated reporter pairs. Paired pulse periods of different reporters are indicated by a grey line. Mean and s.d. are indicated. ns = not significant (paired t-test). nn indicates number of puncta and n indicates number of primordia/embryos from N experiments. **I**, Plot of the normalized temporal correlation coefficients of the indicated fluorescent reporters. Mean and s.d. are indicated. \*\*\*\**P* < 0.0001 (Kruskal-Wallis test). nn indicates number of puncta, n indicates number of primordia/embryos from N experiments.

Since RhoA activates myosin II and myosin II causes F-actin clustering, we further asked whether the formation of active RhoA puncta precedes myosin II clustering, and whether myosin II clustering precedes F-actin clustering. Because addressing these questions requires little interference of signals from neighboring cells, we labeled active RhoA or F-actin in single primordium cells by injecting DNA encoding *cxcr4b:sfGFP-AHPH* or *cxcr4b:F-tractin-mCherry* BAC transgenes into 1-cell stage embryos. Injected DNA is inherited in a mosaic manner and thus labels only a few cells in the primordium (Fig. 4A). The injected embryos were also transgenic for *cxcr4b:sfGFP-AHPH*, *cxcr4b:Myl12.1-mScarlet*, or *cxcr4b:Myl12.1-mNG*. Imaging single cells in such primordia every 2 seconds, confirmed that active RhoA, myosin II clustering and F-actin accumulation are correlated (Fig. 4B–D, H, Video 7). Cross-correlating the fluorescence intensities of the active RhoA, myosin II and F-actin reporters indicates that RhoA activation precedes myosin II clustering and F-actin accumulation by 7 to 13 seconds, respectively, while myosin II clustering and F-actin accumulation cannot temporally be resolved (Fig. 4E–G). Likely due to noise, the lag times between RhoA activation, myosin II clustering and F-actin accumulation are not perfectly additive. However, the lag time between RhoA activation and myosin II activation/F-actin accumulation reflects the signaling sequence of the RhoA pathway and is consistent with observations in other scenarios (Bement et al., 2015; He et al., 2010; Michaux et al., 2018; Munjal et al., 2015; Nishikawa et al., 2017). Thus, RhoA activation leads to actomyosin contraction within a few seconds in the primordium cells — apart from the apical constrictions in the forming neuromasts — mostly on the basal and outer borders of the collective.

**Figure 4.**
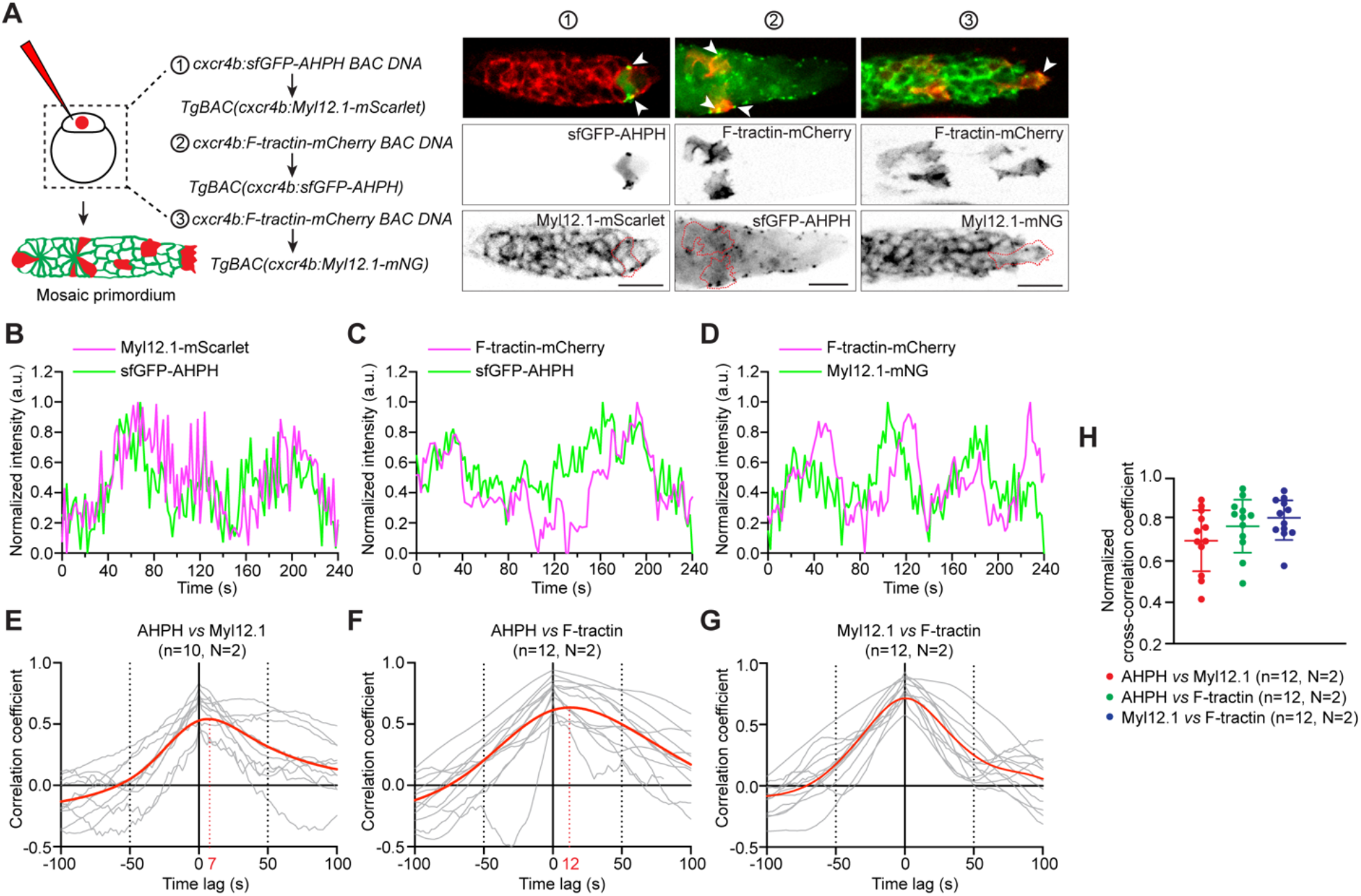
RhoA signaling and actomyosin dynamics are correlated in the cells of the primordium. **A**, Left: Schematic of the strategy to generate mosaic expression of indicated transgenes in the primordium. Right: Images from Video 7 showing the mosaic expression of the indicated transgenes in primordia expressing the indicated reporters. Arrowheads indicate fluorescent reporter clustering and red dotted lines outline the mosaically labelled primordium cells. Images are sum-projected z-stacks. Scale bars, 20 µm. **B**–**D**, Plot of the fluorescent reporter intensities normalized to the mean fluorescent intensity against time in the mosaically labeled primordium cells shown in A. a.u., arbitrary units. **E–G**, Plots of the temporal cross-correlation functions of the fluorescent reporter intensities in the mosaically labeled primordium cells shown in A. Red line is a spline fit to the average of the cross-correlation functions. Lag times are indicated in red on the x-axis. **H**, Plot of normalized temporal cross-correlation coefficients of the fluorescent intensity dynamics between the indicated signals in the mosaically labeled cells shown in A. Mean and s.d. are indicated. n indicates number of cells and N indicate number of experiments.

### Positive and negative feedbacks from the RhoA signaling network tune RhoA activity

Oscillation of RhoA activity has been attributed to a combination of positive and negative feedbacks. RhoA signaling can be amplified by local accumulation of active RhoA through advection (Munjal et al., 2015) or slowed diffusion (Graessl et al., 2017), RhoGEF recruitment by active RhoA (Graessl et al., 2017; Lin et al., 2021) or microtubules (Azoitei et al., 2019; Lin et al., 2021), and stress-induced RhoA activation (Bailles et al., 2019). On the other hand, active RhoA can be turned off by the accumulation of actomyosin (Munjal et al., 2015) and delayed RhoGAP recruitment — often through F-actin — (Graessl et al., 2017; Michaux et al., 2018; Segal et al., 2018), and myosin II-independent ROCK signaling (Ong et al., 2019). To gain insight into the circuitry that regulates active RhoA pulsing in the primordium, we asked how the behavior of active RhoA is affected when we inhibited RhoA, ROCK, or myosin II activity, or F-actin polymerization. Inhibiting RhoA by expressing C3 in the primordium almost completely blocked the formation of active RhoA puncta (Fig. 2C, D). In contrast, blocking ROCK with Rockout or myosin II with blebbistatin for 2 hours did not result in a reduction of the overall intensity of active RhoA in the primordium (Fig. 5B, C, S4I) but increased the number of active RhoA puncta by about 20% (Fig. S4B, S4C) and slowed the migrating primordium (Video 8). Treatment with the inhibitor solvent alone did not affect the puncta number and level of active RhoA and primordium migration (Fig. 5A, S4A, S4I). Contrary to blocking RhoA activity, blocking actin polymerization with Latrunculin A increased the puncta number, size, and intensities of active RhoA (Fig. 5D, S4D, S4I, Video 8), suggesting that F-actin is required to turn RhoA signaling off.

**Figure 5.**
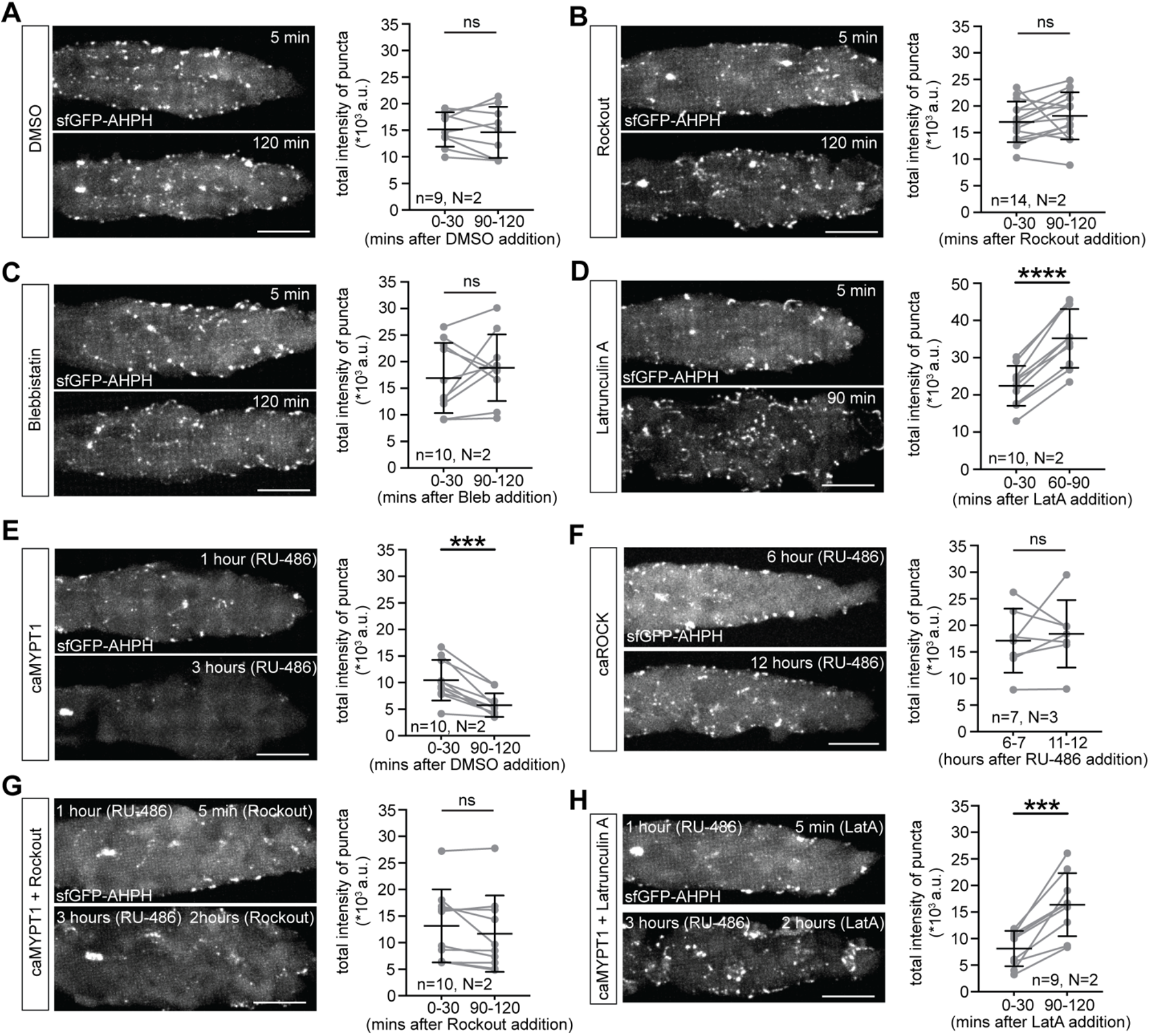
Myosin II and F-actin regulate RhoA activity. **A–D**, Images and quantifications of the total sfGFP-AHPH puncta intensity in the primordium after treatment with DMSO (A), Rockout (B), Blebbistatin (C), and Latrunculin A (D). The time stamp indicates minutes after drug addition (Video 8). Each dot in the quantification represents the average of 7 time points of the indicated time periods. Images were taken every 5 minutes. Mean and s.d. are indicated, and each pair of grey dots connected by a line represents the same embryo. \*\*\*\**P* < 0.0001 (Two-tailed paired t test). ns = not significant. n indicates number of primordia/embryos and N indicates number of experiments. All images are maximum-projected z-stacks. Scale bars, 20 µm. **E**–**F,** Images and quantifications of total sfGFP-AHPH puncta intensity in the primordium expressing caMYPT1 (E) or caROCK (F). The time stamp indicates hours after RU-486 addition (Video 8). Each dot in the quantifications represent the average of 7 (E) and 12 (F) time points of the indicated time periods. Images were taken every 5 minutes. Mean and s.d. are indicated, and each pair of grey dots connected by a line represents the same embryo. ns = not significant and \*\*\**P* = 0.0002 (Two-tailed paired t test). n indicates number of primordia/embryos and N indicates number of experiments. Images are maximum-projected z-stacks. Scale bars, 20 µm. **G**–**H,** Images and quantifications of total sfGFP-AHPH puncta intensity in the primordium expressing caMYPT1 in embryos treated with Rockout (G) or Latrunculin A (H). The time stamp indicates minutes after drug addition (Video 8). caMYPT1 expression in the primordium was induced by adding RU-486 one hour prior to adding the drugs. Each dot in the quantification represents the average of 7 time points of the indicated time periods. Images were taken every 5 minutes. Mean and s.d. are indicated, and each pair of grey dots connected by a line represents the same embryo. ns = not significant and \*\*\**P* = 0.0005 (Two-tailed paired t test). n indicates number of primordia/embryos and N indicates number of experiments. All images are maximum-projected z-stacks. Scale bars, 20 µm.

Because treatment with blebbistatin blocks myosin II activity in the entire embryo, we also blocked myosin II activity specifically in the primordium. For this, we induced caMYPT1 expression in the migrating collective using the *cxcr4b:LexPR-LexOP:caMYPT1* line. Similar to blocking RhoA activity by expressing C3 in the primordium (Fig. 2C, D) — and different from embryos treated with blebbistatin (Fig. 5C), this caused a rapid decrease in RhoA activity to almost undetectable levels within two hours after caMYPT1 induction (Fig. 5E, S4E). Consistent with the reduction in active RhoA, caMYPT1-expressing primordia also ceased to migrate (Videos 4, 8). The difference in reduction of active RhoA in these two scenarios is likely due to the stronger reduction of myosin II activity in caMYPT1-expressing primordia than in primordia of blebbistatin-treated embryos. In control embryos treated with RU-486 but lacking the *cxcr4b:LexPR-LexOP:caMYPT1* transgene, active RhoA level and distribution and primordium migration were unaffected (Videos 4, 8). This indicates that myosin II activity is required for RhoA activation. If the levels of active myosin II are limiting the levels of RhoA activation, then increasing myosin II activity should increase RhoA activity. To test this idea, we induced caROCK expression in the migrating primordium using the *cxcr4b:LexPR-LexOP:caROCK* line. Although expression of caROCK in the primordium increased the intensity of Myl12.1-mScarlet (Fig. S2B, S4K), the intensity of sfGFP-AHPH was unchanged (Fig. 5F, S4F, L). Thus, while myosin II activity is necessary for RhoA activation it is not sufficient to increase active RhoA levels above wild-type levels.

Because ROCK activates myosin II and myosin II activates RhoA in the primordium (Fig. 5B, C, E), reducing ROCK and myosin II activity together should yield the same effect. Contrary to this expectation, we found that Rockout treatment of embryos expressing caMYPT1 in the primordium suppressed the nearly complete inactivation of RhoA observed in untreated embryos expressing caMYPT1 in the primordium (Fig. 5G, S4G, J, Video 8). This suggests that ROCK activity is required for caMYPT1 function, in addition to ROCK’s well documented role in inactivating MYPT1 through phosphorylation on MYPT1’s C-terminal domain (Garrido-Casado et al., 2021; Khromov et al., 2009) which is deleted in caMYPT1 (Smutny et al., 2017). Also, blocking actin polymerization in embryos, in which myosin II function was reduced, should revert the dampening effect of reduced myosin II activity on RhoA activity and increase RhoA activity because active myosin II contracts the F-actin network. Consistent with this reasoning, Latrunculin A-treated embryos expressing caMYPT1 in the primoridum over-activate RhoA signaling to the same degree as Latrunculin A-treated embryos not expressing caMYPT1 (Fig. 5H, S4H, J, Video 8).

Together, these results suggest that myosin II activation is required for RhoA activation and clustering while F-actin is necessary to dampen RhoA activity. Although we cannot exclude actomyosin-independent activation of a diffuse pool of RhoA that was observed by single molecule imaging in early worm embryos (Michaux et al., 2018), our observations are consistent with the ideas that actomyosin contractions concentrate active RhoA (Munjal and Lecuit, 2014) through advection and F-actin aids in the delayed recruitment of RhoGAPs to turn RhoA off (Bement et al., 2015; Graessl et al., 2017; Michaux et al., 2018). These two properties could then constitute the feedforward and feedback loops required for RhoA and myosin II pulsing (Allard and Mogilner, 2013).

### RhoA stimulates actin polymerization at the tip and actin flow at the base of protrusions in the primordium

In many migrating cells, RhoA signaling activates actomyosin to contract and retract the cell’s rear (Parsons et al., 2010; Webb et al., 2002; Yamada and Sixt, 2019). To explore whether RhoA signaling promotes primordium migration in a similar manner, we assessed the distribution of active RhoA and myosin II in single primordium cells. Single cell labeling was achieved by DNA injection of BAC transgenes into one-cell stage embryos, which results in a mosaic inheritance of the DNA in only a few cells during embryonic development. The injected DNA coded for *cxcr4b:sfGFP-AHPH* or *cxcr4b:Myl12.1-mNG* to label active RhoA or myosin II, respectively, and the injected embryos were transgenic for *prim:mem-mCherry* to co-label the membranes of the primordium cells fluorescently. This analysis revealed that active RhoA frequently localizes to the back of cells in the primordium and to the base of their protrusions (Fig. 6A). Myosin II localization is similar to that of active RhoA but myosin II is enriched more at the base of the protrusions than active RhoA (Fig. 6A). Together with the observations that myosin II activity, at least in some instances, contributes to F-actin flow (Lin et al., 1996; Ponti et al., 2004; Wilson et al., 2010; Yolland et al., 2019) and correlates with protrusion retraction (Mishra et al., 2019; Vicente-Manzanares et al., 2011), this suggested that RhoA signaling activates myosin II at the base of protrusions to haul the F-actin network inwards and retract the protrusion of primordium cells. To test this idea, we first asked whether active RhoA and myosin II clustering correlates with protrusion retraction. Imaging the dynamics of active RhoA and myosin II with F-actin in the front cells of the primordium indicates that active RhoA — though not in all cases — and myosin II clusters at the base of protrusions just before protrusions start to retract (Fig. 6B–D). To quantify this correlation, we cross-correlated protrusion dynamics with Myl12.1-mNG or sfGFP-AHPH puncta dynamics in cells at the front of the primordium (Fig. 6E-G). This analysis revealed that protrusive activity is negatively correlated with sfGFP-AHPH and Myl12.1-mNG clustering (Fig. 6G). Importantly, blocking RhoA-mediated signaling by treating embryos with blebbistatin, Rockout, or expressing caMYPT1 in the primordium blocked protrusion retraction (Fig. 6H, Video 9). To assess the consequence of increased RhoA pulsing on protrusion dynamics, we generated mosaic primordia consisting of mostly host wild-type cells and a few donor cells that co-expressed Arhgef11 and H2A-TagBFP upon RU-486 exposure by blastomere transplantation (Fig. 6I). The donor cells were also expressing Myl12.1-mNG and F-tractin-mCherry (Fig. 6J). In Arhgef11-expressing cells, the frequencies of Myl12.1 pulses and protrusion formation were increased (Fig. 6K) but the protrusions were shorter than in wild-type control cells (Fig. 6L), probably because the time to extend protrusions before RhoA-triggered protrusion retraction was reduced. Together these observations indicate that pulses of RhoA-mediated myosin II activation at the base of the protrusion triggers protrusion retraction and set the protrusion-retraction frequency.

**Figure 6.**
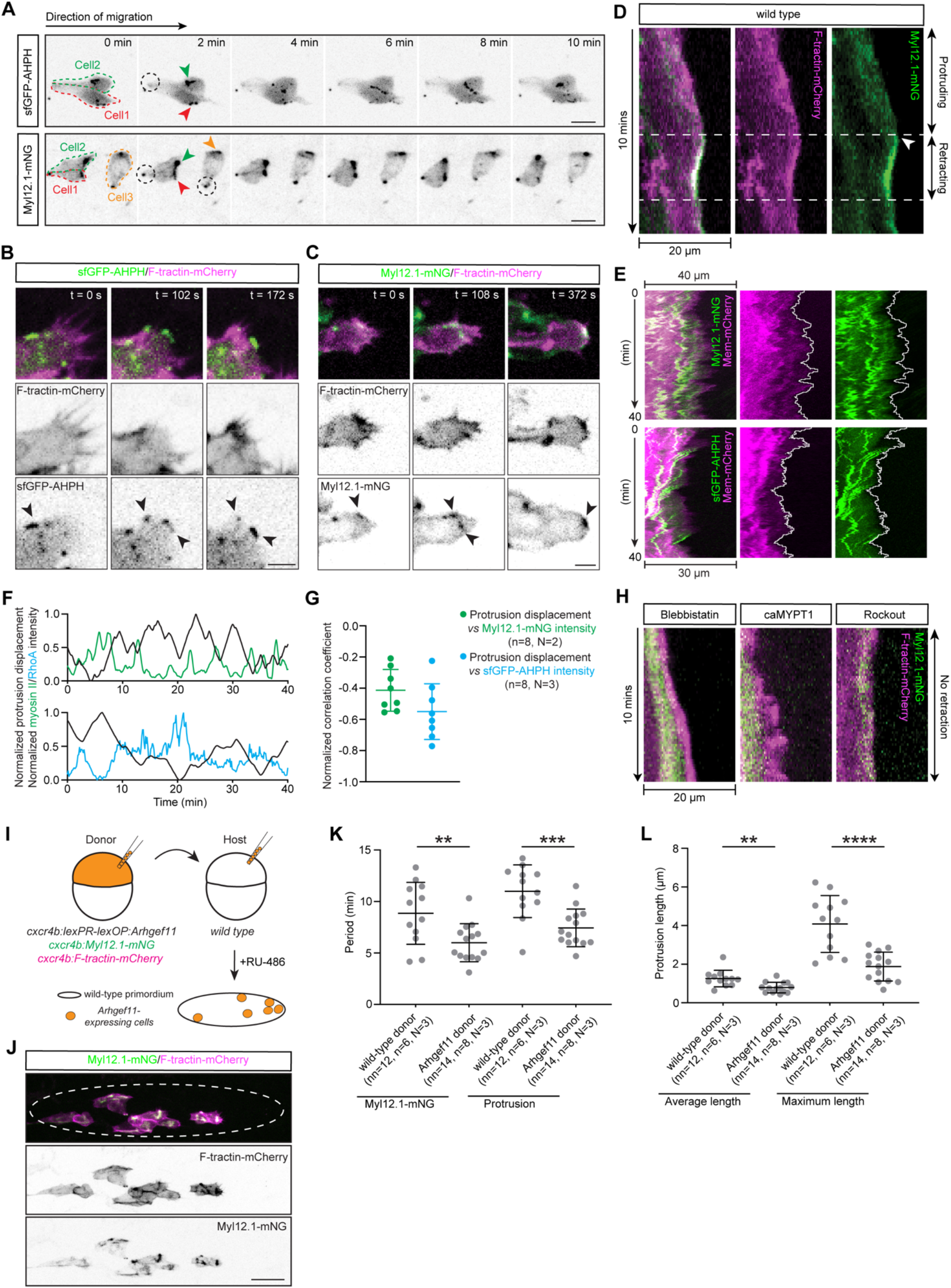
RhoA signaling stimulates protrusion retraction in the primordium cells. **A**, Time series of a few primordium cells labeled with sfGFP-AHPH (top) or Myl12.1-mNG (bottom). Individual cells are outlined by dotted lines. Arrowheads indicate sfGFP-AHPH and Myl12.1-mNG puncta at the base of protrusions in the front of the cells. Dotted black circles indicate sfGFP-AHPH and Myl12.1-mNG puncta at the apical constrictions of the cells. Images are maximum-projected z-stacks. Scale bar, 10 µm. **B**, Time series of F-tractin-mCherry and sfGFP-AHPH localization in a primordium tip cell. Images are maximum-projected z-stacks. Arrowheads indicate the locations of sfGFP-AHPH clustering at the base and tip of protrusions. Scale bar, 5 µm. **C,** Time series of F-tractin-mCherry and Myl12.1-mNG localization in a mosaically labeled tip cell of the primordium. Images are maximum-projected z-stacks. Arrowheads indicate the locations of Myl12.1-mNG clustering at the base of a protrusion. Scale bar, 5 µm. **D**, Kymographs of the F-tractin-mCherry and Myl12.1-mNG fluorescent signals along the protrusion of a mosaically labeled primordium tip cell. Arrowhead indicates the formation of a Myosin II cluster. The two dotted lines indicate the protrusion/retraction transitions. **E–F**, Kymographs (E) of the Myl12.1-mNG (top) or sfGFP-AHPH (bottom) and mem-mCherry fluorescent signals along the protrusions of primordium tip cells in wild-type embryos. The white line traces the tip of the protrusion. The time lapse video was registered to the first apical constriction and spans 40 minutes with a temporal resolution of 12 seconds. The displacement of the protrusion tip normalized to the mean displacement and the intensity of Myl12.1-mNG and sfGFP-AHPH within the protrusion normalized to the mean Myl12.1-mNG and sfGFP-AHPH intensity, respectively, are plotted in F. **G**, Quantifications of the normalized cross-correlation coefficient between the protrusion tip displacement and the intensity of Myl12.1-mNG and sfGFP-AHPH, respectively. n indicates number of primordia/embryos and N indicates number of experiments. **H**, Kymographs of the F-tractin-mCherry and Myl12.1-mNG fluorescent signals along the protrusions of the primordium tip cells in embryos treated with Blebbistatin, Rockout, or expressing caMYPT1 in the primordium. **I**, Experimental design for the generation of chimeric primordia consisting of mostly wild-type cells and a few cells that are labelled with Myl12.1-mNG and F-tractin-mCherry and that will express Arhgef11 upon RU-486 addition. **J,** Representative image of chimeric primordium consisting of mostly wild-type cells (not labelled) and a few Myl12.1-mNG and F-tractin-mCherry-co-labeled cells that are expressing Arhgef11. The dashed line indicates the boundary of the primordium. Scale bar, 20 µm. **K,** Quantifications of the periods for Myl12.1-mNG puncta pulses and protrusions in wild-type cells and Arhgef11-expressing cells placed in wild-type host primordia. nn indicates number of cells, n indicates number of primordia/embryos, N indicates number of experiments. \*\**P* = 0.0067 (Two-tailed unpaired t test), \*\*\**P* = 0.0004 (Two-tailed unpaired t test). **L,** Quantifications of the average and maximum protrusion lengths in wild-type cells and donor Arhgef11-expressing cells placed in wild-type primordia. The average and maximum protrusion lengths are measured by tracking the dynamics of donor cell protrusion displacement in videos spanning 40 minutes with a temporal resolution of 12 seconds. nn indicates number of cells, n indicates number of primordia/embryos, N indicates number of experiments. \*\**P* = 0.0024 (Two-tailed unpaired t test), \*\*\*\**P* < 0.0001 (Two-tailed unpaired t test).

Next, we tested whether the myosin II activity is essential for retrograde F-actin flow in the protrusions of primordium cells. For this, we tracked retrograde F-actin flow in protrusions of the primordium’s front cells in embryos treated with blebbistatin or expressing caMYPT1 in the primordium. To monitor inhibition of myosin II clustering, these embryos also expressed Myl12.1-mNG in the primordium. While myosin II clustered at the base of protrusions and F-actin flowed rearwards in the cells of control embryos (Fig. 7A–C, Video 9), we detected little to no F-actin flow in embryos with blocked myosin II activity (Fig. 7A–C, Video 9). Similarly, blocking ROCK activity with Rockout also blocked myosin II clustering and resulted in stalled F-actin flow in the protrusions (Fig. 7A, B, Video 9). In contrast to blocking myosin II activation, however, blocking ROCK activity additionally impaired F-actin polymerization at the tip of the primordium cell protrusions (Fig. 7A, D, E, Video 9). Consistent with this observation, we also detected small clusters of active RhoA in some but not all protrusion tips (Fig. 6B) — possibly because some clusters of active RhoA are below the detection limit of confocal microscopy. Together, these observations indicate that RhoA signaling promotes primordium migration in two ways. First, it activates myosin II at the base of protrusions to initiate protrusion retraction and generate rear-directed F-actin flow. Second, RhoA signaling induces F-actin polymerization at the tip of the cell protrusions in the primordium through ROCK but independent of myosin II activity, possibly by activating LIM kinase and Diaphanous — two pathways that promote actin polymerization (Bros et al., 2019; Lee et al., 2015; Maekawa et al., 1999; Watanabe et al., 1997).

**Figure 7.**
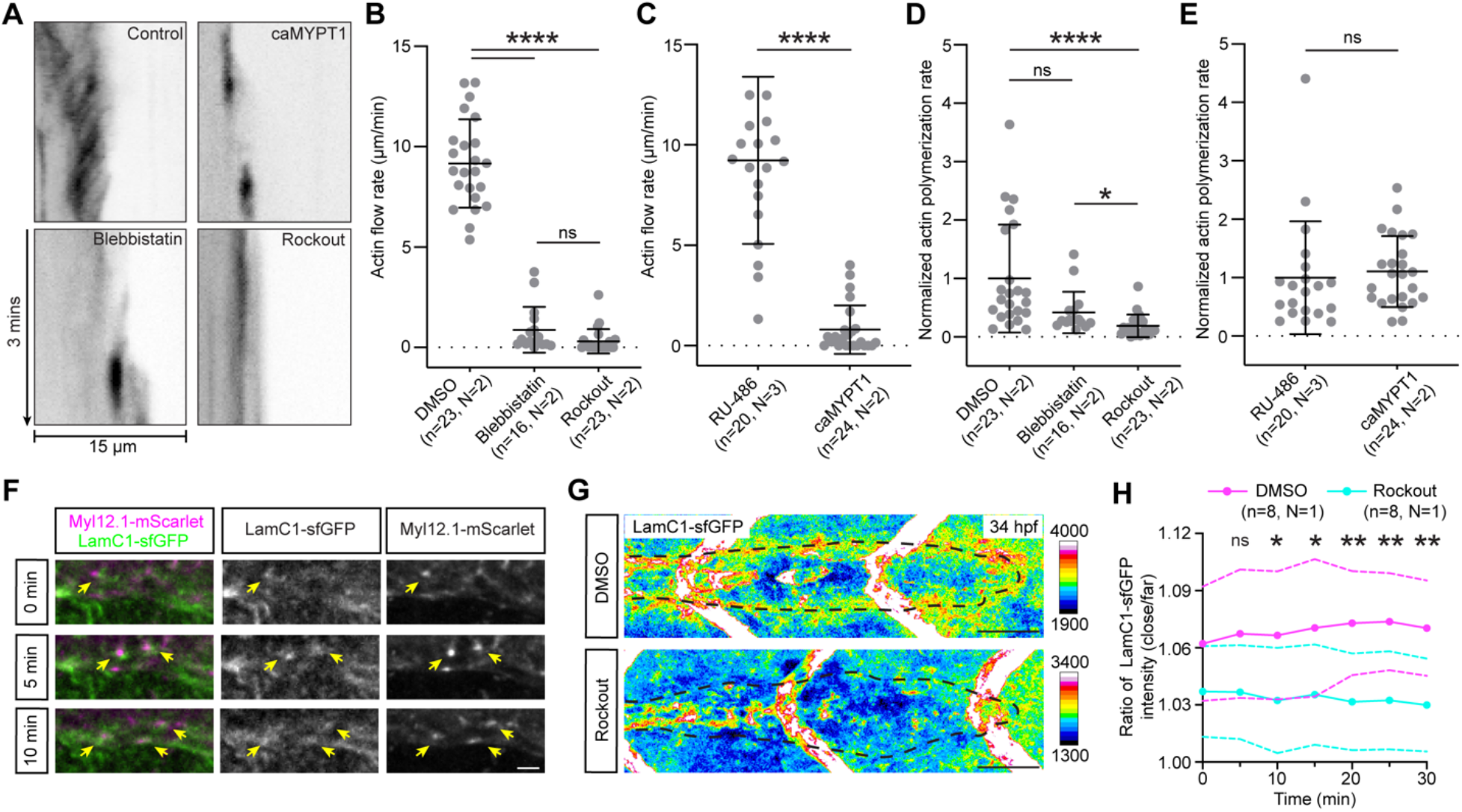
RhoA signaling stimulates actin polymerization and actin flow in the protrusions of primordium cells and induces traction against the extracellular matrix. **A**, Kymographs of the F-tractin-mCherry signal along the protrusions of primordium tip cells in embryos treated with DMSO, blebbistatin, or Rockout, or expressing caMYPT1 (Video 9). **B–C**, Quantification of the actin flow rate in the tip of primordium cells of embryos treated with DMSO, blebbistatin, or Rockout (B) and in the tip of primordium cells of wild-type control embryos and embryos expressing caMYPT1 in the primordium (C). Mean and s.d. are indicated. Each dot represents the actin flow rate measured in a single primordium cell. \*\*\*\**P* < 0.0001 (Mann-Whitney test for Control (RU-486) versus caMYPT1 and ordinary one-way ANOVA for control (DMSO) versus blebbistatin versus Rockout). n indicates number of primordia/embryos and N indicates number of experiments. **D–E**, Quantification of the actin polymerization rate in the tip of primordium cells of embryos treated with DMSO, blebbistatin, or Rockout (D) and in the tip of primordium cells of wild-type embryos and embryos expressing caMYPT1 in the primordium (E). Mean and s.d. are indicated. Each dot represents the actin polymerization rate measured in a single primordium cell and is normalized to the mean of control condition in each group. D: \**P* = 0.0454, \*\*\*\**P* < 0.0001 (Unpaired test for Control (RU-486) versus caMYPT1 and Kruskal-Wallis for Control (DMSO) versus blebbistatin versus Rockout); ns = not significant. n indicates number of primordia/embryos and N indicates number of experiments. **F**, Still images from a time series of Myl12.1-mScarlet and LamC1–sfGFP localization at the basal sides of primordium cells. Images are maximum-projected z-stacks. Arrows indicate transient adjacent Myl12.1-mScarlet and LamC1-sfGFP clusters. Scale bar, 5 µm. **G**, Distribution of LamC1-sfGFP around the primordium outlined by a dotted line in 34 hpf embryos treated with DMSO (top) or Rockout (bottom). LamC1-sfGFP fluorescence intensity is pseudo-colored as a heat map. Images are maximum-projected z-stacks. Note scale of fluorescence intensity is different between control-treated and Rockout-treated embryos. Scale bar, 25 µm. **H**, Time series of the ratio of the mean LamC1–sfGFP fluorescence intensities in two elliptical annuli that are close to and distant from the border of the primordium for embryos treated with DMSO (pink) or Rockout (cyan), respectively (Video 10). The two elliptical annuli were defined using the border of the primordium as a reference. The close elliptical annulus starts 1.6 um inside the the border of the primordium and extends 1.6 um outwards, and the distant elliptical annulus starts 4.8 um from the primordium’s border and extends to 8.0 um outwards. Z-stacks were collected every 5 minutes. Solid curves indicate the mean and dotted curves indicate the s.d. ns = not significant (t = 5 min), \**P* = 0.0436 (t = 10 min), \**P* = 0.0438 (t = 15 min), \*\**P* = 0.0074 (t = 20 min), \*\**P* = 0.0061 (t = 25 min), \*\**P* = 0.0056 (t = 30 min) (Multiple unpaired t tests). n indicates number of primordia/embryos and N indicates number of experiments.

### The primordium generates traction on the basement membrane through RhoA signaling

The primordium migrates on top of a basement membrane that it deforms as it pushes itself forward (Metcalfe, 1985; Yamaguchi et al., 2022). Together with our observation that the primordium needs RhoA signaling for motility, this suggests that RhoA-mediated rear-directed F-actin flow — coupled to the outside through friction — should be required for traction generation and basement membrane deformation. To test this idea, we first asked whether myosin II clustering correlated with basement membrane deformation. As observed in other contexts (Hagedorn et al., 2013; Harris et al., 1980), basement membrane deformations cause the basement membrane to buckle and wrinkle (Yamaguchi et al., 2022). To assess wrinkling, we labeled the basement membrane in embryos using GFP-tagged Laminin-γ1 (LamC1) — a core basement membrane component (Pastor-Pareja, 2020) — expressed under the control of its endogenous genomic architecture on a BAC. The embryos also expressed mScarlet-tagged Myl12.1 in the primordium to assess myosin II localization. In such embryos, myosin II clusters formed and disassembled in the primordium next to transient LamC1-sfGFP aggregation in the basement membrane (Fig. 7F). Second, we asked whether basement membrane wrinkling required RhoA signaling. In contrast to LamC1-sfGFP aggregation around the circumference of the primordium in control embryos (Fig. 7G, H, Video 10), such aggregates were not detectable around primordia in embryos treated with the ROCK inhibitor Rockout (Fig. 7G, H, Video 10). Together, these observations are consistent with the idea that RhoA-mediated actomyosin contractions are coupled to the basement membrane and cause the basement membrane to buckle as the primordium pushed itself forward.

In summary, our study provides two major insights into the force generating machinery in a migrating tissue. First, we find that appropriate levels of RhoA signaling are required for primordium motility. This is different from RhoA’s role in collective migration in other contexts where loss of RhoA activity results in failure of cells to rearrange (Pirraglia et al., 2013; Xu et al., 2008; Xu et al., 2011), to properly adhere to each other (Bastock and Strutt, 2007; Lebreton and Casanova, 2013), or to polarize along the front-rear axis (Matthews et al., 2008), and is more similar RhoA’s requirement in single cell motility — though RhoA is required for rear retraction rather than protrusion dynamics in this context, indicating that the primordium uses RhoA for motility rather than tissue cohesion. Second, we show that RhoA signaling stimulates primordium motility through two distinct processes: at the tip of protrusions, RhoA signaling promotes actin polymerization, and at the base of protrusions, RhoA signaling activates myosin II to pull the F-actin network rearward and retract protrusions. These roles for RhoA that also have been recently identified in cultured cells (Azoitei et al., 2019; Hu et al., 2022; Lee et al., 2015; Machacek et al., 2009; Martin et al., 2016; Pertz et al., 2006; Tkachenko et al., 2011; Yang et al., 2015). Together, these RhoA activities generate rearward directed actin flow that is then transmitted — partly through integrins (Yamaguchi et al., 2022) — to the substrate and propels the primordium forward. Since many cells move in groups, it is likely that this force generating machinery underlies the motility of collectively migrating cells in other contexts.

## Materials and Methods

### Zebrafish husbandry

This study was performed in accordance with recommendations in the Guide for the Care and Use of Laboratory Animals of the National Institutes of Health. Zebrafish husbandry and experimental procedures were approved by the NYU Grossman School of Medicine Institutional Animal Care and Use Committee (IACUC), under protocols: IA16-00788_AMEND202100320.

### Zebrafish strains

Embryos were raised at 28.5 °C and staged as previously described (Kimmel et al., 1995). The following transgenic lines have previously been reported: *Tg(prim:mem-mCherry)* (Wang et al., 2018), *Tg(cldnB:lyn_2_GFP)* (Haas and Gilmour, 2006), *TgBAC(lamC1:lamC1-sfGFP)* (Yamaguchi et al., 2022), *TgBAC(cxcr4b:F-tractin-mCherry)* (Yamaguchi et al., 2022), and *TgBAC(cxcr4b:Myl12.1-mScarlet)* (Yamaguchi et al., 2022).

### Generation of transgenic lines

To express genes and reporters using the *cxcr4b* regulatory region, we used the BAC clone DKEY-169F10, which was obtained from ImaGenes GmbH, Germany. DKEY-169F10 contains a 69 kb genomic DNA fragment that spans the entire *cxcr4b* locus. As previously described, we modified this BAC clone to include a transgenesis marker (either *cryaa:dsRed* or *my17:mScarlet*) and *tol2* sequences in the BAC backbone (Fuentes et al., 2016). Using these intermediate BAC clones, we used galK-mediated BAC recombineering to generate the final transgenic BAC clones (Warming et al., 2005). All modified BAC clones were characterized by EcoRI restriction digest and by sequencing a PCR amplicon of the modified region. Each final BAC clone was purified with Nucleobond BAC 100 Kit (Clonetech) and co-injected with 40 ng/ul *tol2* mRNA into one-cell stage embryos. Stable transgenic lines were established by out-crossing the adult founder fish injected with the transgene and raising the embryos that express the transgenesis marker and — if visible — the protein of interest in the primordium.

#### TgBAC(cxcr4b:PAK-PBD-sfGFP)

The active-Rac/Cdc42 localization reporter sequence PAK-PBD was previously characterized (Manser et al., 1994; Srinivasan et al., 2003). We identified the zebrafish *pak1 PAK-PBD* sequence by aligning it with the human PAK1 PAK-PBD. The zebrafish *pak1 PAK-PBD* sequence (amino acids 64-149 in Pak1) together with four amino acid linker sequence (amino acids: LSGR) was amplified from zebrafish cDNA by PCR and cloned into the targeting vector for BAC recombineering. A cassette consisting of *PAK-PBD -sfGFP-FRT-galK-FRT*, flanked by homology arms 583 bp upstream from the beginning of *cxcr4b* exon 2 and 433 bp downstream of the *cxcr4b* stop codon, respectively, was inserted to replace the *cxcr4b* exon 2 coding sequence (amino acids 6-358). This transgene expresses the first five amino acids from *cxcr4b* exon 1 fused to *PAK-PBD* from the *cxcr4b* promoter. The full name of this transgenic line is *TgBAC(cxcr4b: PAK-PBD-sfGFP)-p1* and the transgenesis marker is *cryaa:dsRed*.

#### TgBAC(cxcr4b:sfGFP-AHPH)

The AHPH domain of anillin was previously characterized and used as a localization reporter for active RhoA (Piekny and Glotzer, 2008; Priya et al., 2015; Sun et al., 2015; Tse et al., 2012). The zebrafish anillin AHPH domain sequence (amino acids 678–1171 of Anillin isoform 1) together with a ten amino acid linker sequence (amino acids: SGLRSRAQAS) was amplified from zebrafish cDNA by PCR and cloned into a targeting vector. The full name of this transgenic line is *TgBAC(cxcr4b:sfGFP-AHPH)-p1* and the transgenesis marker is *cryaa:dsRed*.

#### TgBAC(cxcr4b:Myl12.1-mNeonGreen)

The *myl12.1* coding sequence was amplified from the targeting plasmid *pUC19-homology-arm-myl12.1-mScarlet-FRT-galK-FRT-homology-arm* (Yamaguchi et al., 2022) and the mNeonGreen coding sequence was amplified from *mNeonGreen-mTurquoise2* plasmid (Addgene #98886) (Mastop et al., 2017). The full name of this transgenic line is *TgBAC(cxcr4b:Myl12.1-mNeonGreen)-p3* and the transgenesis marker is *myl7:mScarlet*.

#### TgBAC(cxcr4b:LexPR-LexOP:C3-T2A-H2A-Cerulean)

To generate the *cxcr4b:LexPR-LexOP:C3-T2A-H2A-Cerulean* transgene, we used the modified BAC clone DKEY-169F10 which carries the transgenesis marker *myl7:mScarlet* and modified it further in a two-step manner. First, a cassette consisting of *LexPR-FRT-galK-FRT* was inserted into the BAC as described above. Second, a cassette containing *LexOP-C3-T2A-H2A-Cerulean-SV40polyA-loxP-Amp^R^-loxP*, flanked 5’ with 318 bp and 3’with 362 bp homology arms, was inserted in the BAC about 15 kb downstream of the stop codon of *cxcr4b* by ampicillin-mediated selection. Note that Amp^R^ cassette was not removed from the transgene. The coding sequence of zebrafish codon optimized *C3* transferase was obtained from IDT. The LexPR and LexOP sequences were amplified from the *mitfa-LexPR-P2A-Cerulean* plasmid and the *pCrysb-ECFP-LexOP-mCherry-NrasQ61K* plasmid (Kenyon et al., 2018), respectively, both were kind gifts from Tatjana Sauka-Spengler. Founder fish were additionally tested for H2A-Cerulean expression after RU-486 treatment. The full name of this transgenic line is *TgBAC(cxcr4b:LexPR-LexOP:C3-T2A-H2A-Cerulean)-p1*.

#### TgBAC(cxcr4b:LexPR-LexOP:C3Δ12-T2A-H2A-Cerulean)

To generate the *TgBAC(cxcr4b:LexPR-LexOP-*C3Δ12*-T2A-H2A-Cerulean)* control strain, we mutated the coding sequence of C3 by the CRISPR-Cas9 based gene-editing. Briefly, tracrRNA and crRNA pair targeting the coding sequence of C3 (5’-TGGATCGATATAACCACCCT-3’) was designed and purchased from IDT. Purified protein Cas9-NLS was purchased from QB3 MacroLab at UC Berkeley (https://macrolab.qb3.berkeley.edu/). The injection mix containing Cas9-NLS protein, crRNA and tracrRNA was heat-activated and injected into one-cell stage *TgBAC(cxcr4b:LexPR-LexOP:C3-T2A-H2A-Cerulean)-p1* embryos. The stable transgenic line was established by out-crossing adult fish injected with the CRISPR/Cas9 mix as embryos and raising the embryos with no C3 activity after RU-486 administration. The 12 bp deletion in the C3 coding sequence was verified by PCR and sequencing. The full name of this transgenic line is *TgBAC(cxcr4b:LexPR-LexOP:*C3Δ12*-T2A-H2A-Cerulean)-p1*.

#### TgBAC(cxcr4b:LexPR-LexOP:caMYPT1-T2A-H2A-TagBFP)

To generate the *cxcr4b:LexPR-LexOP:caMYPT1-T2A-H2A-TagBFP* transgene, we used the intermediate BAC clone DKEY-169F10 with the transgenesis marker *myl7:mScarlet* and the insertion of *LexPR* sequence into the *cxcr4b* coding sequence that is described above. We constructed caMYPT1 by amplifying the truncated coding sequence of zebrafish protein phosphatase regulatory subunit 12A which encodes N-terminal of Ppplr12A (amino acids 1–301) from cDNA (Smutny et al., 2017). The cassette containing *LexOP-caMYPT1-T2A-H2A-TagBFP-SV40polyA-loxP-Amp^R^-loxP* was then inserted into the BAC as described above for *TgBAC(cxcr4b:LexPR-LexOP:C3-T2A-H2A-Cerulean)*. Injected embryos were raised to adulthood and outcrossed. Founder fish were additionally tested for H2A-TagBFP expression after RU-486 treatment. The full name of this transgenic line is *TgBAC(cxcr4b:LexPR-LexOP:caMYPT1-T2A-H2A-TagBFP)-p1*.

#### TgBAC(cxcr4b:LexPR-LexOP:caROCK-T2A-H2A-TagBFP)

To generate the *cxcr4b:LexPR-LexOP:caROCK-T2A-H2A-TagBFP* transgene, we used the above-described modified BAC clone DKEY-169F10 with the transgenesis marker *myl7:mScarlet* and the insertion of *LexPR* sequence into the *cxcr4b* coding sequence. To construct the caROCK, the coding sequence without the pleckstrin homology (PH) domains (Amano et al., 1999) for zebrafish *Rock2a* (amino acid sequence of Rock2a 1–1033, accession number: NP_777288) was amplified from zebrafish maternal cDNA by PCR and assembled into the following targeting cassette in a *pUC19* plasmid: *LexOP-caROCK-T2A-H2A-TagBFP-SV40polyA-loxP-Amp^R^-loxP*. Founder fish were additionally tested for H2A-TagBFP expression after RU-486 treatment. The full name of this transgenic line is *TgBAC(cxcr4b:LexPR-LexOP:caROCK-T2A-H2A-TagBFP)*.

#### TgBAC(cxcr4b:LexPR-LexOP:Arhgef11-T2A-H2A-TagBFP)

To generate the *cxcr4b:LexPR-LexOP:Arhgef11-T2A-H2A-TagBFP* transgene, we used the above-described modified BAC clone DKEY-169F10 with the transgenesis marker *myl7:mScarlet* and the insertion of *LexPR* sequence into the *cxcr4b* coding sequence. The coding sequence for zebrafish *Arhgef11*, a homolog of *rhoGEF2* in fruit fly and *ARHGEF11* in humans (Panizzi et al., 2007), was amplified from zebrafish 36 hpf cDNA by PCR and assembled into the following targeting cassette in a *pUC19* plasmid: *LexOP-Arhgef11-T2A-H2A-TagBFP-SV40polyA-loxP-Amp^R^-loxP*. Founder fish were additionally tested for H2A-TagBFP expression after RU-486 treatment. For the reported experiment, we used double transgenic embryos of the two lines we established. The full names of the transgenic lines are *TgBAC(cxcr4b:LexPR-LexOP:Arhgef11-T2A-H2A-TagBFP)-p1* and *TgBAC(cxcr4b:LexPR-LexOP-Arhgef11-T2A-H2A-TagBFP)-p2*.

### Generation of chimeric primordium

For chimeric analysis, we transplanted 20-50 cells from donor embryos at ∼1000 cell stage into host embryos of the same stage. All donor embryos were transgenic for *<u>TgBAC</u>*(*cxcr4b:H2A-GFP*) and were either wild type or had one of the following transgenes: *TgBAC(cxcr4b:LexPR-LexOP:C3-T2A-H2A-Cerulean)* or *TgBAC(cxcr4b:LexPR-LexOP:Arhgef11-T2A-H2A-TagBFP)*. Host embryos were either wild type or transgenic for *TgBAC*(*cxcr4b:H2A-mCherry*). At ∼28 hpf, we treated the embryos with 5 μM RU-486 and the drug was present throughout imaging. Chimeric primordia with the donor cells expressing *cxcr4b:H2A-GFP* were identified with a Zeiss Axio Zoom V16 stereo microscope and were imaged on an inverted Nikon W1 spinning disk confocal to record time lapse videos. To confirm whether the chimeric primordium was positive for C3 expression, imaged embryos were digested and the genomic DNA was extracted to perform PCR amplification of C3. To confirm whether the chimeric primordium was positive for Arhgef11 expression, embryos were checked for TagBFP expression on an upright Leica SP8 confocal microscope.

### Generation of mosaic primordium

To generate mosaic primordia, we co-injected 1 nl BAC DNA coding for the transgene of interest (about 50 ng/µl) with *tol2* mRNA (40 ng/µl) into one-cell-stage embryos that also were transgenic for different RhoA signaling reporters as indicated in the figures. Injected embryos with mosaic transgene expression in the primordium were then selected for imaging.

### Whole mount immunofluorescent staining

To stain against phospho-Myosin Light Chain 2 (Thr18/Ser19), we fixed *TgBAC(cxcr4b:Myl12.1-mNeonGreen)-p3* embryos at 31 hpf with 4% PFA/PBST for 2 hours at room temperature, dehydrated them in a methanol series, and stored the embryos in 100% methanol (Millipore-Sigma, cat no. 322415-100ML) overnight at - 20°C. Embryos were rehydrated using a series of 50% and 30% methanol in PBST and blocked in 1% BSA/PBST for 1 hour at room temperature. The embryos were incubated in rabbit anti-phospho-Myosin Light Chain 2 (Thr18/Ser19) antibody (1:200, catalog # 3678, Cell Signaling Technology) in 1% BSA/PBST overnight at 4°C. Embryos were washed four times with PBST and incubated with goat anti-rabbit Alexa568 secondary antibody (1:500, Invitrogen, catalog # A-11011) in PBST overnight at 4°C. Embryos were washed four times with PBST and incubated with donkey anti-goat Cy3 tertiary antibody (1:500, Jackson ImmunoResearch, catalog # 705-165-147) in PBST overnight at 4°C. Embryos were washed four times with PBST, then mounted in 0.5% low melt agarose/Ringer’s solution for imaging.

### Pharmacological treatments

To assess the migration distance of the primordium, embryos of different genetic backgrounds [(1) *cldnB:lyn_2_GFP*, (2) *TgBAC(cxcr4b:LexPR-LexOP:C3-T2A-H2A-Cerulean)*; *cldnB:lyn_2_GFP*, (3) *TgBAC(cxcr4b:LexPR-LexOP:C3Δ12-T2A-H2A-Cerulean)*; *cldnB:lyn_2_GFP*, (4) *TgBAC(cxcr4b:LexPR-LexOP:caMYPT1-T2A-H2A-TagBFP)*; *cldnB:lyn_2_GFP*, (5) *TgBAC(cxcr4b:LexPR-LexOP:caROCK-T2A-H2A-TagBFP)*; *cldnB:lyn_2_GFP*, (6) *TgBAC(cxcr4b:LexPR-LexOP:Arhgef11-T2A-H2A-TagBFP)*] and control sibling embryos were dechorionated at 24 hpf and incubated in fish water with either RU-486 (final concentration 1 μM, Millipore Sigma, catalog # M8046) or ethanol until 48 hpf. To assess the migration distance of the primordium in embryos treated with different small molecular inhibitors, *cldnB:lyn_2_GFP* embryos were dechorionated at 24 hpf and incubated in the fish water supplemented with either Rockout (final concentration 50 μM, Millipore Sigma, catalog # 555553), blebbistatin (final concentration 1 μM, Millipore Sigma, catalog # B0560) or 1% DMSO solvent control until 48 hpf.

To perturb downstream components of RhoA/ROCK signaling for live-imaging of active-RhoA, Latrunculin A (final concentration 2 μM; Abcam, catalog # ab144290), blebbistatin (final concentration 100 μM; Millipore Sigma, catalog # B0560), and Rockout (final concentration 200 μM, Millipore Sigma, catalog # 555553) diluted in 1 mL of Ringer’s solution was added at the beginning of the imaging. To overexpress caMYPT1, we first incubated 32 hpf *TgBAC(cxcr4b:sfGFP-AHPH)*; *Tg(prim:mem-mCherry)*; *TgBAC(cxcr4b:LexPR-LexOP:caMYPT1-T2A-H2A-TagBFP)* embryos with RU-486 (final concentration 5 μM, Millipore Sigma, catalog # M8046) for 1 hour before we imaged the embryos in Ringer’s solution supplemented with Rockout, Latrunculin A or 1% DMSO solvent control as described above. RU-486 was added from 34 hpf onwards. To over express caROCK, we first incubated 24 hpf *TgBAC(cxcr4b:sfGFP-AHPH)*; *TgBAC(cxcr4b:Myl12.1-mScarlet)*; *TgBAC(cxcr4b:LexPR-LexOP:caMYPT1-T2A-H2A-TagBFP)* embryos with RU-486 (final concentration 5 μM, Millipore Sigma, catalog # M8046) for 6 hours before imaging the embryos in Ringer’s solution. For long-time live-imaging of *TgBAC(cxcr4b:sfGFP-AHPH)*; *TgBAC(cxcr4b:Myl12.1-mScarlet)* embryos with the treatment of Rockout and blebbistatin, we used a final concentration of 200 μM for both Rockout and blebbistatin and added the inhibitors at the beginning of the imaging.

### Confocal imaging

Confocal imaging was performed either on an upright Leica SP8 confocal microscope equipped with a HC PL APO 40x/1.10 W CORR CS2 Leica (catalog # 15506357, Fig. 1A-C, Fig. S1A, B, Fig. 2C, Fig. S2A-D), or a HC PL Fluotar 20x/0.50 Leica (catalog # 15506503, Fig. 2E, Fig. S2E) objective, an inverted Nikon W1 spinning disk confocal equipped with an Apo LWD 40x/NA 1.15 Nikon (catalog # MRD77410, Fig. 4A, Fig. 5A-E, G-H, Fig. 6A-D, H, Fig. 7F, G), a Plan Fluor 40x/NA1.3 Nikon (catalog # MRH01401, Fig. 2H, Fig. 5F), or an Apo 60x/NA1.40 Nikon (catalog # MRD71600, Fig. 3A-D, Fig. S3B, C, Fig. 6E, J, Fig. 7A) objective. For imaging with the Leica SP8 confocal microscope, embryos were mounted in 0.5% low-melt agarose (National Diagnostics, EC-205)/Ringer’s solution supplemented with 0.4 mg/mL MS-222 anesthetic on a coverslip (Fig. 1A-C, Fig. S1A, B, Fig. 2C, Fig. S2A-D) or a plastic dish (Fig. 2E, Fig. S2E). Embryos were kept at 28 °C with a heated stage on the Leica SP8 confocal microscope (Warner Instruments, Quick Exchange Heated Base, QE-1) for long-time live imaging (Fig. 2E, Fig. S2E). The laser power was calibrated using a power meter (X-Cite Power Meter Model, Lumen Dynamics, XR2100). The HyD detectors and the photon-counting mode were used for the acquisition of the fluorescent intensity. For imaging with the Nikon W1 spinning disk confocal microscope, embryos were mounted in 0.5% low-melt agarose (National Diagnostics, EC-205) in Ringer’s solution supplemented with 0.4 mg/ml MS-222 anesthetic on a glass-bottom dish (Fig. 2H, Fig. 3A-D, Fig. S3B, C, Fig. 4A, Fig. 5A-H, Fig. S4, Fig. 6A-E, H, J, Fig. 7A, F, G) and were kept at 30 °C using a heated chamber (Tokai Hit incubation system STXG-TIZWX-SET). The two-channel sequential scan setting was used to prevent fluorescence bleed-through.

To assess the effects of overexpression of C3, caMYPT1, caROCK, and Arhgef11 on the reporters for active RhoA (sfGFP-AHPH, Fig. 2C, Fig. S2C) and myosin II (Myl12.1-mNG or Myl12.1-mScarlet, Fig. S2A, B, D), embryos were imaged on the Leica SP8 confocal microscope at 33 hpf (C3, caMYPT1, and Arhgef11) or 43 hpf (caROCK). For overexpression of C3, embryos were treated with 5 µM of RU-486 from 27 hpf onwards. For overexpression of ca-MYPT1, embryos were treated with 5 µM of RU-486 from 30 hpf onwards. For overexpression of caROCK and Arhgef11, embryos were treated with 5 µM of RU-486 from 24 hpf onwards.

For long-term live imaging of primordium migration upon induction of C3, caMYPT1, caROCK, or Arhgef11 expression, *cldnB:lyn_2_GFP* or *prim:mem-mCherry* embryos that were also transgenic for *TgBAC(cxcr4b:LexPR-LexOP:C3-T2A-H2A-Cerulean)*, *TgBAC(cxcr4b:LexPR-LexOP:caMYPT1-T2A-H2A-TagBFP), TgBAC(cxcr4b:LexPR-LexOP:caROCK-T2A-H2A-TagBFP), or TgBAC(cxcr4b:LexPR-LexOP:Arhgef11-T2A-H2A-TagBFP)* were imaged on the Leica SP8 confocal microscope. Embryos were mounted on a plastic dish. Live imaging was performed with a 20x/NA 0.5 Leica objective every 20 minutes for a least 8 hours. The embryos were treated with 5 µM of RU-486 for the indicated times (Fig. 2E, Fig. S2E) before the start of the imaging.

To image chimeric embryos with nuclei-labelled primordium cells that were either wild-type or transgenic for *TgBAC(cxcr4b:LexPR-LexOP:C3-T2A-H2A-Cerulean)* or *TgBAC(cxcr4b:LexPR-LexOP:Arhgef11-T2A-H2A-TagBFP)* (Fig. 2H), we treated the embryos with 5 µM of RU-486 at ∼28 hpf, and started imaging after 4 hours of RU-486 treatment. Imaging was performed on an inverted Nikon W1 spinning disk confocal at 30 °C. Chimeric embryos were imaged for at least 5 hours. The time interval was set to 7 minutes for chimeric embryos that were transgenic for *TgBAC(cxcr4b:LexPR-LexOP:C3-T2A-H2A-Cerulean)* or 5 minutes for wild-type embryos and embryos that were transgenic for *TgBAC(cxcr4b:LexPR-LexOP:Arhgef11-T2A-H2A-TagBFP)*.

To perform imaging with high spatial-temporal resolution for quantifying puncta intensity dynamics at the outer border cells (Fig. 3A-D, Fig. S3B, C) and the tip cells (Fig. 6E) of the primordium, embryos were imaged every 12 seconds for 40 minutes within a 12 µm depth centered around the middle plane of the outer border and tip cells. Z-step size was set to 1 μm.

To image puncta intensity dynamics for mosaically labelled cells, embryos were imaged every 2 seconds for 10 mins within an 11 µm depth centered around the middle plane of interest (Fig. 4A), or every 10 seconds for 10 mins within a 30 µm depth (Fig. 6A-D, H). Z-step size was set to 1 μm. To perform imaging for actin retrograde flow measurements (Fig. 7A), embryos were imaged every 2 seconds for 3 mins within a 3 µm depth centered around the middle plane of interest. Z-step size was set to 1 µm.

To image the effects of small molecule inhibitors against ROCK, myosin II, and F-actin, expression of caMYPT1, or expression of caROCK on active RhoA using the sfGFP-AHPH reporter, embryos of the appropriate genotype were imaged every 5 mins from 33 hpf for 2 hours or from 32 hpf for 6 hours for the expression of caROCK with a Z-step size of 1 μm (Fig. 5A-H, Fig. S4). After the first one or two time points, we added the small molecule inhibitors directly to the medium in the dish in which the embryos were mounted. To induce expression of caMYPT1, we first incubated 32 hpf *TgBAC(cxcr4b:sfGFP-AHPH)*; *Tg(prim:mem-mCherry)*; *TgBAC(cxcr4b:LexPR-LexOP:caMYPT1-T2A-H2A-TagBFP)* embryos with RU-486 for 1 hour before imaging with or without the treatment of Rockout or Latrunculin A. To induce expression of caROCK, we first incubated 26 hpf *TgBAC(cxcr4b:sfGFP-AHPH)*; *TgBAC(cxcr4b:Myl12.1-mScarlet)*; *TgBAC(cxcr4b:LexPR-LexOP:caROCK-T2A-H2A-TagBFP)* embryos with RU-486 for 6 hour before imaging.

To image the accumulation of LamC1-sfGFP under the primordium expressing the myosin II reporter (Fig. 7F), embryos were imaged at 31 hpf. The Z-step size was set to 0.4 μm and Z-stacks were acquired every 5 minutes for 30 minutes. To image the accumulation of LamC1-sfGFP under the primordium with the inhibition of ROCK (Fig. 7G), embryos were treated with either 200 µM Rockout (Millipore Sigma, catalog # 555553) or 1% DMSO solvent control at 30 hpf. Imaging was performed 2 hours after treatment. The Z-step size was set to 1 μm and Z-stacks were acquired every 5 minutes for 30 minutes.

### Image analysis and quantification

All image analyses were performed in Fiji and MATLAB 2021a (Mathworks) with custom-written macros and scripts.

#### Localization analysis of sfGFP-AHPH, Myl12.1-mNG, and F-tractin in the primordium

The maximum-intensity projection of sfGFP-AHPH and Myl12.1-mNG images were cropped and a distance of 100 µm from the tip of the primordium was defined as the migrating part for quantification. To analyze the sfGFP-AHPH localization, the sfGFP-AHPH channel was sum-projected and converted into a binary mask. The mask was denoised with cycles of voxel erosion and dilation. To analyze the signal of active-RhoA in the primordium, the mean intensity of the sfGFP-AHPH signal was obtained in the masked area. We then subtracted 1.25*mean intensity from the projected image. We further created a ring-like binary mask that is 4.2 µm in width and is between 4.2 – 8.4 µm from the periphery of the primordium. The mask was then applied to the subtracted image and the mean fluorescence intensity was obtained. The image created with a peripheral mask was resliced from the tip of the primordium and the fluorescence intensity profile from the tip to 100 µm away was obtained. Analysis of the localization of Myl12.1-mNG and F-tractin in the primordium was the same with sfGFP-AHPH except that for Myl12.1-mNG, we generated the binary mask with the membrane-mCherry channel and we did not subtract 1.25*mean intensity from the image to include all the signal intensity in the analysis.

#### Measuring puncta number, size, and intensity of sfGFP-AHPH and Myl12.1-mNG

The maximum-intensity projection of sfGFP-AHPH and Myl12.1-mNG images were cropped and a distance of 100 µm from the tip of the primordium was defined as the migrating part for the quantification. First, a binary mask was created by thresholding the sum-projected membrane-mCherry intensity signal to include the primordium. This mask was then applied to the maximum-projected sfGFP-AHPH or Myl12.1-mNG Z-stacks to obtain the mean intensity of sfGFP-AHPH and Myl12.1-mNG in the primordium. Then, the standard deviation plus the mean intensity was subtracted and the image was binarized for counting the number and size of puncta. The number and size of puncta was counted using the Analyze Particle function in Fiji with the size set to 0.1 μm^2-Infinite for sfGFP-AHPH and Myl12.1-mNG. The binarized region of interest (ROI) of all the puncta was then applied to the maximum-projected Myl12.1-mNG image to get the mean intensity of each single puncta. For measuring puncta properties in a time series movie, the movie was first registered using a custom written macro. The registered movie was then used and quantified as described above.

#### Migration distance measurement

To quantify the cumulative migration distance of the primordium, the Z-stacks from the time-lapse image sequences were maximum-intensity projected and the tip of the primordium was tracked using the Manual Tracking plugin provided by Fabrice Cordelieres in Fiji (https://imagej.nih.gov/ij/plugins/manual-tracking.html). The kymograph was generated using the maximum-intensity projected Z-stack sequences with the KymoResliceWide plugin by Eugene Katrukha and Laurie Young (https://imagej.net/KymoResliceWide). To quantify the migration ratio of the primordium at 48 hpf in the different scenarios (genetic and pharmacological perturbations), images were taken with a Leica 165M FC Fluorescent Stereo Microscope equipped with a Leica DFC345 FX camera. We then measured the distances from the ear to the tip of the tail and from the ear to the tip of the primordium using Fiji.

#### Tracking of the cell nuclei in the chimeric primordium

To track the 3D position of cell nuclei in the chimeric primordium, we imported the time lapse videos of chimeric primordium with host cells labeled with H2A-mCherry and donor cells labelled with H2A-GFP into Imaris Version 10.0 (Bitplane, Oxford Instruments). The spot tool in the Imaris software was used as described previously (Colak-Champollion et al., 2019). The tracks that were wrongly produced in the Imaris were manually corrected. The tracked data was then imported into MATLAB for the analyses of *speed*, *neighbor-neighbor distances*, and *directional index* as described previously (Colak-Champollion et al., 2019).

#### Puncta intensity dynamics measurement

To quantify the intensity dynamics of AHPH, Myl12.1, and F-tractin puncta at the border of the primordium, maximum-intensity projections of the two-channel videos were first made and registered using a custom written macro. A binary mask was then created using one of the channels of the registered videos. The mask was then applied to the maximum intensity projected Z-stacks to obtain the mean intensity of each channel. The mean intensity was subtracted from each channel to obtain the final Z-stacks for kymograph. For the analysis of primordia co-labelled with sfGFP-AHPH and Myl12.1-mScarlet, the AHPH channel was used to create the binary mask; For the analysis of primordium co-labelled with sfGFP-AHPH and F-tractin-mCherry or Myl12.1-mNG and F-tractin-mCherry, the F-tractin-mCherry channel was used to create the binary mask and after subtracting the mean intensity of each channel, masks were created from the final sfGFP-AHPH and Myl12.1-mNG maximum intensity projected Z-stacks and were applied to the F-tractin-mCherry channel to obtain the signals that are co-localized with sfGFP-AHPH and Myl12.1-mNG, respectively. For the analysis of primordia co-labelled with Myl12.1-mNG and mem-mCherry, the mem-mCherry channel was used to create the binary mask. To generate kymographs for each fluorescent intensity channel from the two borders of each primordium, we used a 10 μm-wide and 70-μm long line ROI along the front-back margin (Fig. S3A). The kymograph was then generated using the KymoResliceWide plugin from E. Katrukha and L. Young (https://imagej.net/KymoResliceWide). A 2-4 μm-wide segmented line ROI was defined by manually tracing a given puncta over time in first fluorescent intensity channel. The ROI was applied to the second fluorescent intensity channel and the fluorescent intensities for each fluorescent channel were extracted along this ROI. For the analysis of primordium co-labelled with sfGFP-AHPH and Myl12.1-mScarlet or Myl12.1-mScarlet and F-tractin-mCherry, kymograph from the Myl12.1-mScarlet channel was used to define the segmented line ROI; For the analysis of primordium co-labelled with sfGFP-AHPH and F-tractin-mCherry, kymograph from the sfGFP-AHPH channel was used to define the segmented line ROI; The fluorescent intensity profiles of each punctate were then imported into a custom-written MATLAB (MathWorks) script for further analysis.

To obtain the cross-correlation and delay time between two different fluorescent signals in mosaically labeled cells (Fig. 4), we first quantified the intensity over time of the two signals. To do that, we created a binary mask from the intensity image of the mosaically expressed transgene and applied this mask to the intensity image of the stably expressed transgene. The resulting intensity image of the stably expressed transgene was used together with the intensity image of the mosaically expressed transgene and we manually selected an ROI and quantified the intensity over time. A rectangular ROI was manually drawn to include area with visible puncta. After extracting the intensity profile of these two signals, we imported the intensity profiles into a custom-written MATLAB script for further analysis.

#### Protrusion dynamics measurement

To correlate the protrusion dynamics and the activity of RhoA and myosin II in the tip cells of the primordium (Fig. 6F-G), we used embryos co-labelled with sfGFP-AHPH and prim:mem-mCherry or Myl12.1-mNG and prim:mem-mCherry. For transplantation experiments (Fig. 6I-J), wild-type donor cells and donor cells expressing Arhgef11 were labelled with Myl12.1-mNG and F-tractin-mCherry. The videos were first registered using the apical constriction points closest to the tip cells or in the transplanted donor cells. A rectangular ROI was then manually drawn to cover cell protrusion within the 40-minute video and the kymograph was generated using the KymoResliceWide plugin from E. Katrukha and L. Young (https://imagej.net/KymoResliceWide) for either both the membrane channel and the sfGFP-AHPH and Myl12.1-mNG channel in tip cells, respectively, or the Myl12.1-mNG and F-tractin-mCherry channels in transplanted donor cells. The resultant kymograph of the membrane channel or the F-tractin-mCherry channel were used to create a mask for the edge of the protrusion. A 1-pixel width ROI was then manually drawn along the edge of the mask. The x coordinates of the edge were used as the displacement of the protrusion. To extract the intensity of sfGFP-AHPH and Myl12.1-mNG along the edge of the protrusion, the 1-pixel width ROI was expanded to 10-pixel and moved horizontally towards the base of the protrusion to cover the sfGFP-AHPH and Myl12.1-mNG signals, respectively. A custom-written macro was used to extract the intensity of sfGFP-AHPH and Myl12.1-mNG within the 10-pixel width ROI in the kymograph. The protrusion edge displacement and the intensity of sfGFP-AHPH and Myl12.1-mNG were then imported into MATLAB for correlation analyses as described below. We used *findpeaks* function in the MATLAB to determine the magnitude of the protrusion edge displacement that we defined as protrusion length in Fig. 6L.

#### Correlation analyses

Fluorescence intensity profiles of puncta across the primordium’s outer border and in mosaically labelled cells were obtained as described above. The fluorescent intensity profiles were imported into a custom-written MATLAB script. The normalized correlation coefficients for the cross-correlation of the two intensity profiles were calculated using the *crosscorr* function. The cross-correlation analyses also provide the delay time between two signals in mosaically labelled cells. A spline fit was then applied to the combined single-ROI cross-correlation functions as shown in Fig. 4E-G. To calculate the pulse period of puncta, auto-correlation analysis was performed. Briefly, we firstly computed the autocorrelation of each intensity profile such that it is unity at zero lag. To find the peaks in the autocorrelation curve, we used *findpeaks* function and define a minimum peak height (*MinPeakheight*) of zero and a minimum peak prominence (*MinPeakProminence*) of 0.05. Distance in time between any two of the closest peaks were averaged to obtain the pulse period of the puncta.

#### Quantification of actin retrograde flow and actin polymerization rates

To quantify the F-actin retrograde flow, a 1-pixel-wide 15 µm ROI line was drawn manually from the center of the cell outwards across the protrusion of a given cell. The kymograph was then generated using the KymoResliceWide plugin. The front of the actin flow was visually identified on the kymograph and manually traced by drawing a line along the front of the actin flow. The actin flow rate was then calculated by measuring the slope of the trace line. To quantify the actin polymerization rate, we used the same kymographs that we used for the F-actin flow quantification. The actin polymerization event was manually identified on the kymograph and a vertical 5 pixel-wide ROI was drawn on the kymograph such that the ROI encompassed the actin polymerization event. The intensity profile of the ROI on the kymograph was then extracted. To calculate the actin polymerization rate, a linear line was fitted to the intensity increase part of the intensity profile using linear regression (Microsoft Excel). The slope of the fitted line was used as the actin polymerization rate.

#### LamC1-sfGFP intensity measurements

The quantification of LamC1-sfGFP intensity due to basement membrane wrinkles has been described previously (Yamaguchi et al., 2022). Briefly, we quantified the intensity of LamC1-sfGFP around the periphery of the primordium (I-close) and the intensity of LamC1-sfGFP beyond the primordium (I-far). We then used the ratio of I-close to I-far as an indication of basement membrane wrinkling. To quantify I-close, we created a 10 pixel-wide (3.25 µm) ring-like binary mask using the primordium membrane signal from the *Tg(prim:mem-mCherry)* transgene as the reference. The LamC1–sfGFP signal at the myotendinous junctions was threshold to create a binary mask and was subtracted from the ring-like binary mask. The final ring-like binary mask was then applied to the LamC1-sfGFP image to extract the average intensity defined as I-close. To quantify I-far, we created a 10-pixel-wide (3.25 µm) ring-like binary mask excluding the myotendinous junction area but extending to 6.5 µm away from the border of the primordium. Mean intensity of the LamC1-sfGFP in the ring area that was 6.5 µm away from the border of the primordium was quantified as the I-far.

#### Statistics

All statistical tests were performed using Prism 9 (GraphPad software). The Kolmogorov– Smirnov (KS) test was used to examine the normality of the data. If the data passed the normality test, the ordinary one-way analysis of variance (ANOVA) test was used for data consisting of more than two groups followed by multiple comparisons as indicated. The two-tailed t-test was used for data consisting of two groups. If the data did not pass the normality test, the Kruskal-Wallis test was used for data consisting of more than two groups followed by multiple comparisons as indicated. The Mann–Whitney test was used for data consisting of two groups.

## Acknowledgements

We thank C. O’keeffe, J. He, B. Zhao, P. Vagni, and J. Torres-Vázquez for critical comments; J. Proietti, S. Pirani, A, Adams, and T. Dunlap for excellent fish care; Y. Deng for advice on microscopy; Y. Bernadskaya and L. Christiaen for the use of Imaris. The use of Leica TCS SP8 confocal (grant S10 RR024708) and the Nikon W1 spinning disk (grant P01A1080192) from NYU Langone’s Microscopy Laboratory (RRID: SCR_017934) is gratefully acknowledged. The NYU Langone’s Microscopy Laboratory is partially supported by Cancer Center Support Grant P30CA016087. This work was supported by the U.S. National Institutions of Health R01NS119449 (H.K.), by NYSTEM training grant C322560GG (W.Q.) and C322560GG (N.Y.), by an American Heart Association postdoctoral fellowship 903886 (W.Q.), by an American Heart Association predoctoral fellowship 20PRE35180164 (N.Y.), and by an NYU Dean’s Undergraduate Research Fund (P. Lis).

## Author contributions

Conceptualization, W.Q., N.Y., and H.K.; Methodology, W.Q., N.Y., M.C., and H.K.; Experiments, W.Q., N.Y., P.L., and H.K.; Data analysis, W.Q., and N.Y.; Writing, W.Q., N.Y., and H.K.; Supervision, H.K.

---

**Figure S1.**
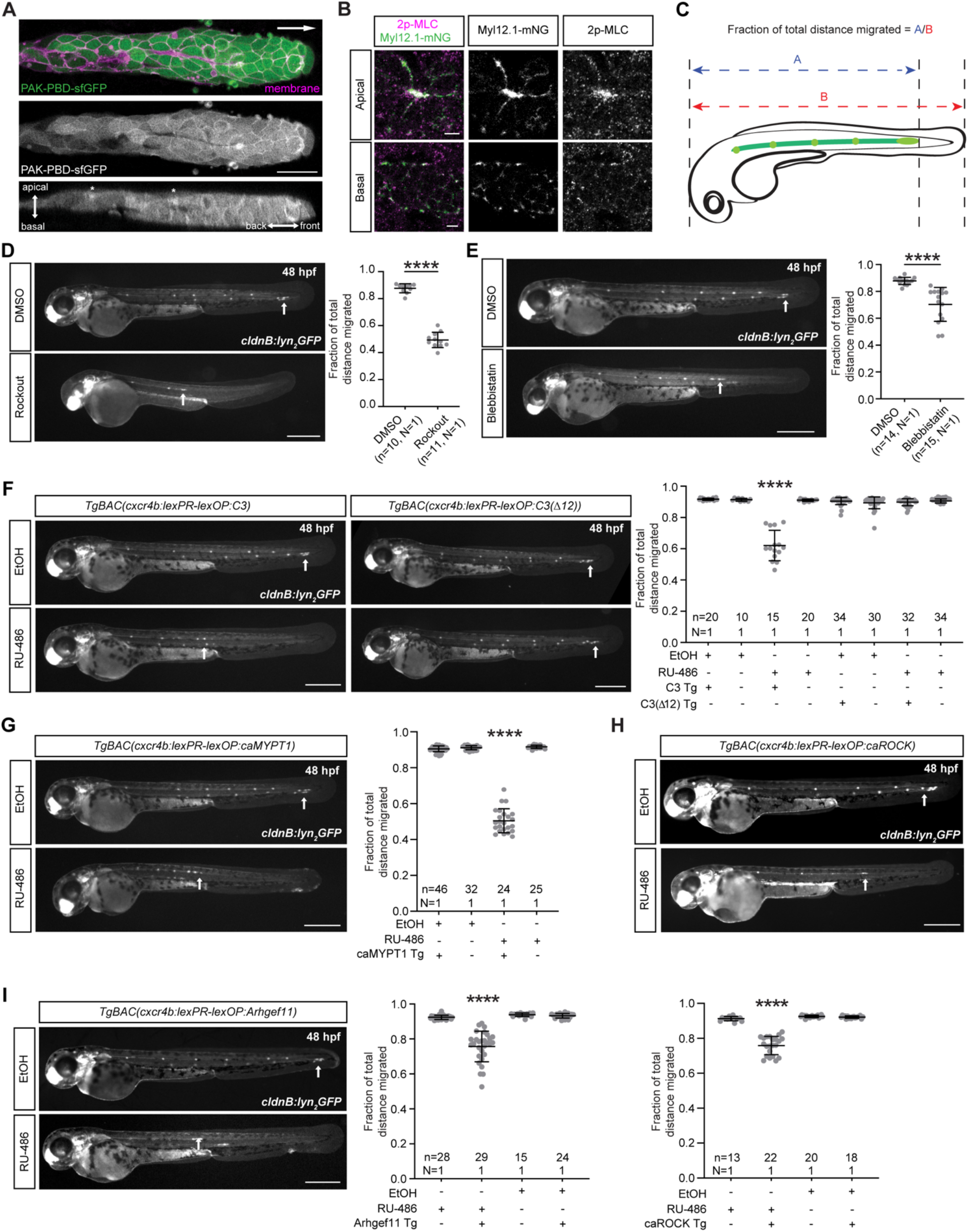
Rac reporter localization in the primordium and migration distance of the primordium in embryos with compromised RhoA-mediated signaling. **A**, Representative image showing the distribution of PAK-PBD-sfGFP in the primordium. Top and middle panels are single xy-sections and bottom panels are single xz-sections through the middle of the primordium from a z-stack. Arrow in top panel indicates direction of primordium migration. Asterisks in lower panel indicate the location of the apical constrictions. Scale bar, 20 µm. **B**, Immunofluorescence staining against phospho-Myosin Light Chain 2 (Thr18/Ser19) in a *cxcr4b:Myl12.1-mNG* embryo. Images are single xy-sections from a z-stack of a forming neuromast in the back of the primordium at the apical and basal sides of the primordium. Scale bars, 5 µm. **C,** Schematic illustration of the quantification of the relative primordium migration distance. **D**, **E**, Representative images and quantifications of primordium migration distances at indicated time points in embryos treated with Rockout (D) or blebbistatin (E). Data represent mean ± s.d.. \*\*\*\**P* < 0.0001 (Unpaired t test). Scale bar, 100 µm. Arrows indicate the primordia’s positions. **F**, Representative images and quantifications of primordium migration distances at indicated time points in embryos treated with RU-486 or vehicle (EtOH) only. Genotypes are indicated. C3(Δ12) is an in-frame deletion in C3 that abrogates C3’s activity. Data represent mean ± s.d.. \*\*\*\**P* < 0.0001 (Kruskal-Wallis test). Scale bar, 100 µm. Arrows indicate the primordia’s positions. **G**, Representative images and quantifications of primordium migration distances at indicated time points in embryos of the indicated genotype and treated with RU-486 or vehicle (EtOH) only. Data represent mean ± s.d.. \*\*\*\**P* < 0.0001 (Kruskal-Wallis test). Scale bar, 100 µm. Arrows indicate the primordia’s positions. **H**, Representative images and quantifications of primordium migration distances at indicated time points in embryos of indicated genotype and treated with RU-486 or vehicle (EtOH) only. Data represent mean ± s.d.. \*\*\*\**P* < 0.0001 (Kruskal-Wallis test). Scale bar, 100 µm. Arrows indicate the primordia’s positions. **I**, Representative images and quantifications of primordium migration distances at indicated time points in embryos of indicated genotype and treated with RU-486 or vehicle (EtOH) only. Data represent mean ± s.d.. \*\*\*\**P* < 0.0001 (Kruskal-Wallis test). Scale bar, 100 µm. Arrows indicate the primordia’s positions. In all panels, n indicates number of primordia/embryos, N indicates number of experiments.

**Figure S2.**
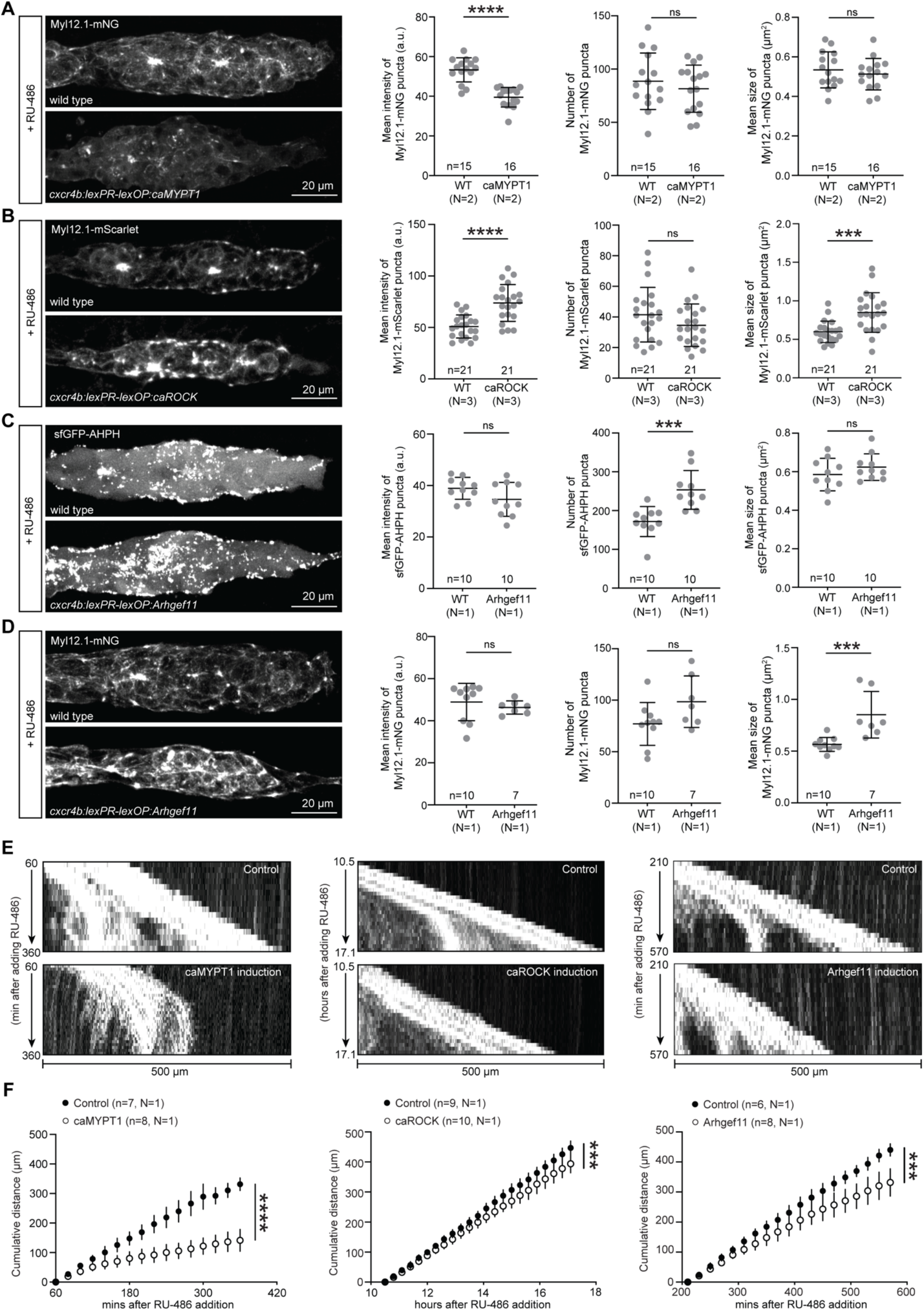
Active RhoA and Myosin II localization and primordium migration in embryos with altered RhoA signaling in the primordium. **A**, Left: Representative images of the Myl12.1-mNG localization pattern in primordia of control embryos (top) and embryos expressing caMYPT1 in the primordium (bottom). Images are maximum-projected z-stacks. Scale bar, 20 µm. Right: Quantification of the mean fluorescence intensity, number, and mean size of Myl12.1-mNG puncta shown in A. Mean and s.d. are indicated. \*\*\*\**P* < 0.0001 (Unpaired t test). **B**, Left: Representative images of the Myl12.1-mScarlet localization pattern in primordia of control embryos (top) and embryos expressing caROCK in the primordium (bottom). Images are maximum-projected z-stacks. Scale bar, 20 µm. Right: Quantification of the mean fluorescence intensity, number, and mean size of Myl12.1-mScarlet puncta shown in B. Mean and s.d. are indicated. \*\*\**P =* 0.0003 and \*\*\*\**P* < 0.0001 (Unpaired t test). **C, D**, Left: Representative images of the sfGFP-AHPH (C) and Myl12.1-mNG (D) localization pattern in primordia of control embryos (top) and embryos expressing Arhgef11 in the primordium (bottom). Images are maximum-projected z-stacks. Scale bar, 20 µm. Right: Quantification of the mean fluorescence intensity, number, and mean size of sfGFP-AHPH (C) and Myl12.1-mNG (D) puncta shown in C and D. Mean and s.d. are indicated. In C, \*\*\**P =* 0.0007 and in D, \*\*\**P =* 0.0004 (Unpaired t test). **E**, Kymograph of primordia in control embryos (top) and embryos expressing caMYPT1 (left), caROCK (middle), or Arhgef11 (right) in the primordium (bottom). The kymographs trace the tip of the primordium shown in Video 4. **F**, Plot of the cumulative primordium migration distance in control embryos (black circles) and embryos expressing caMYPT1 (left), caROCK (middle), or Arhgef11 (right) in the primordium (white circles). Mean and s.d. are indicated. In F, ****P (control vs caMYPT1) < 0.0001, ***P (control vs caROCK) = 0.0005, and ***P (control vs Arhgef11) = 0.0002 (Unpaired t test). In all panels, n indicates number of primordia/embryos, N indicates number of experiments.

**Figure S3.**
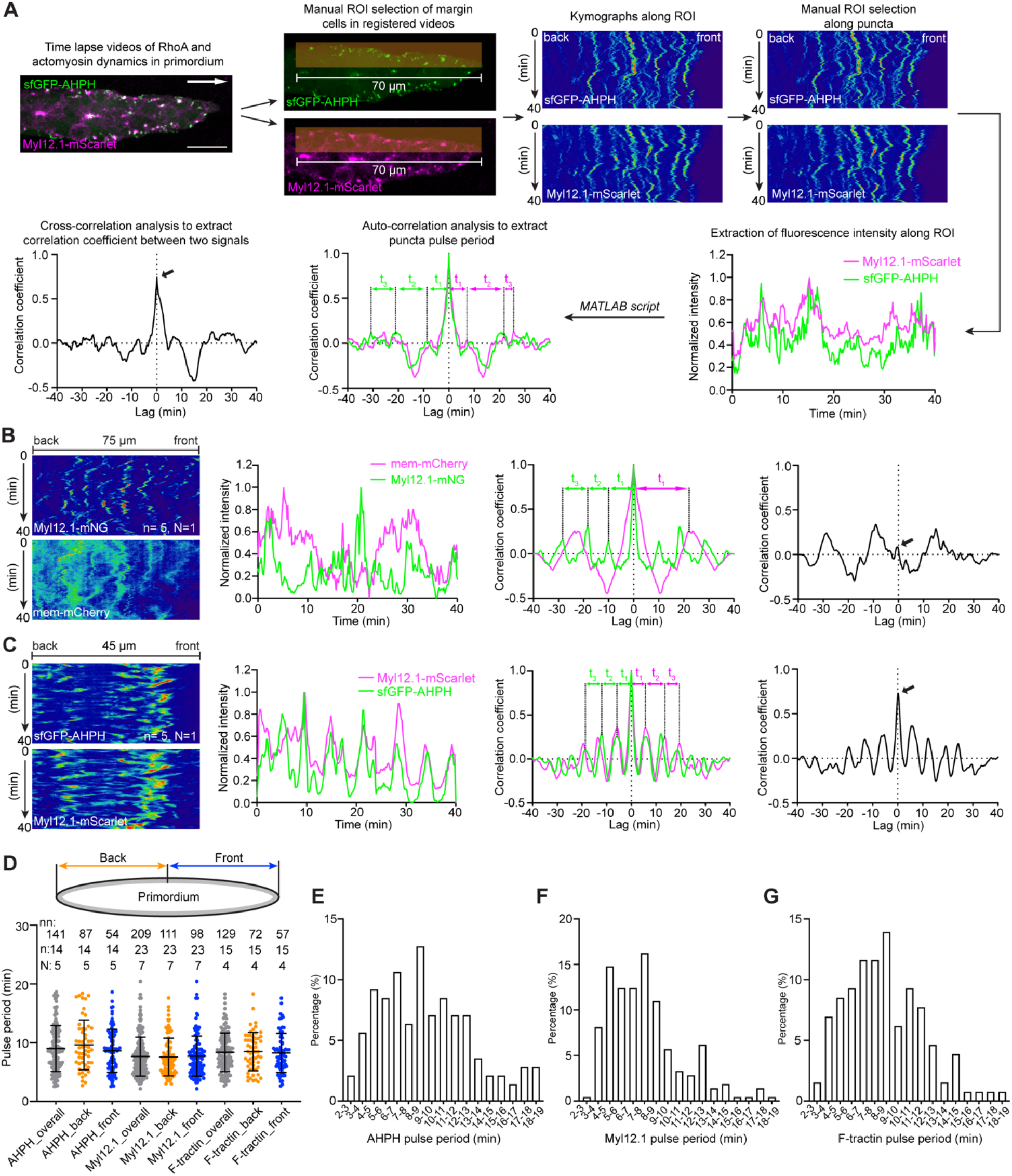
Analysis of the pulsing of RhoA signaling components in the primordium. **A**, Summary of workflow used to analyze the dynamics of AHPH, Myl12.1 and F-tractin reporters in the cells at the margin of the primordium. Maximum-projected z-stacks of 40-min time lapse videos were registered to the first apical constriction (asterisk) and a 10 μm-wide and 70-μm long line ROI along the front-back margin of the primordium was manually defined to generate kymographs for each fluorescent intensity channel. A 2–4 μm-wide segmented line ROI was defined by manually tracing a given puncta over time in the fluorescent intensity image of AHPH-sfGFP or Myl12.1-mNG. The ROI was applied to the other fluorescent intensity channel and the fluorescent intensities for each fluorescent channel were extracted along this ROI. The fluorescent intensity profiles were imported into a custom-written MATLAB script and the normalized correlation coefficients for the auto-correlation and the cross-correlation of the two intensity profiles and the puncta pulse periods were calculated. White arrow in the maximum-projected z-stacks indicates direction of migration, scale bar is 20 μm. Double sided arrows in the auto-correlation graph indicate the puncta pulse periods, which were averaged to obtain the mean pulse period for a single punctate. Black arrow in the cross-correlation graph indicates the largest cross-correlation coefficient for the representative sfGFP-AHPH and Myl12.1-mScarlet intensity profiles. **B–C**, Representative workflows showing the auto-correlation and the cross-correlation analysis of a wild-type primordium expressing Myl12.1-mNG and mem-mCherry (B) and a Arhgef11-overexpressing primordium also expressing sfGFP-AHPH and Myl121-mScarlet (C). Black arrows in the cross-correlation graph indicate the largest cross-correlation coefficients for the representative intensity profiles. n indicates number of primordia/embryos, N indicates number of experiments. **D**, Plot of pooled pulse periods for AHPH, Myl12.1, and F-tractin. The pulse periods were divided into “front” and “back” groups based on the puncta locations along the front-back axis of the primordium’s margin. No significant difference of the pulse periods was observed for puncta at different locations along the margin of the primordium. nn indicates number of the analyzed puncta, n indicated number of analyzed primordia, and N indicates number of experiments. **E–G**, Histograms of binned pulse periods for AHPH (E), Myl12.1 (F), and F-tractin (G) puncta along the margin of the primordium.

**Figure S4.**
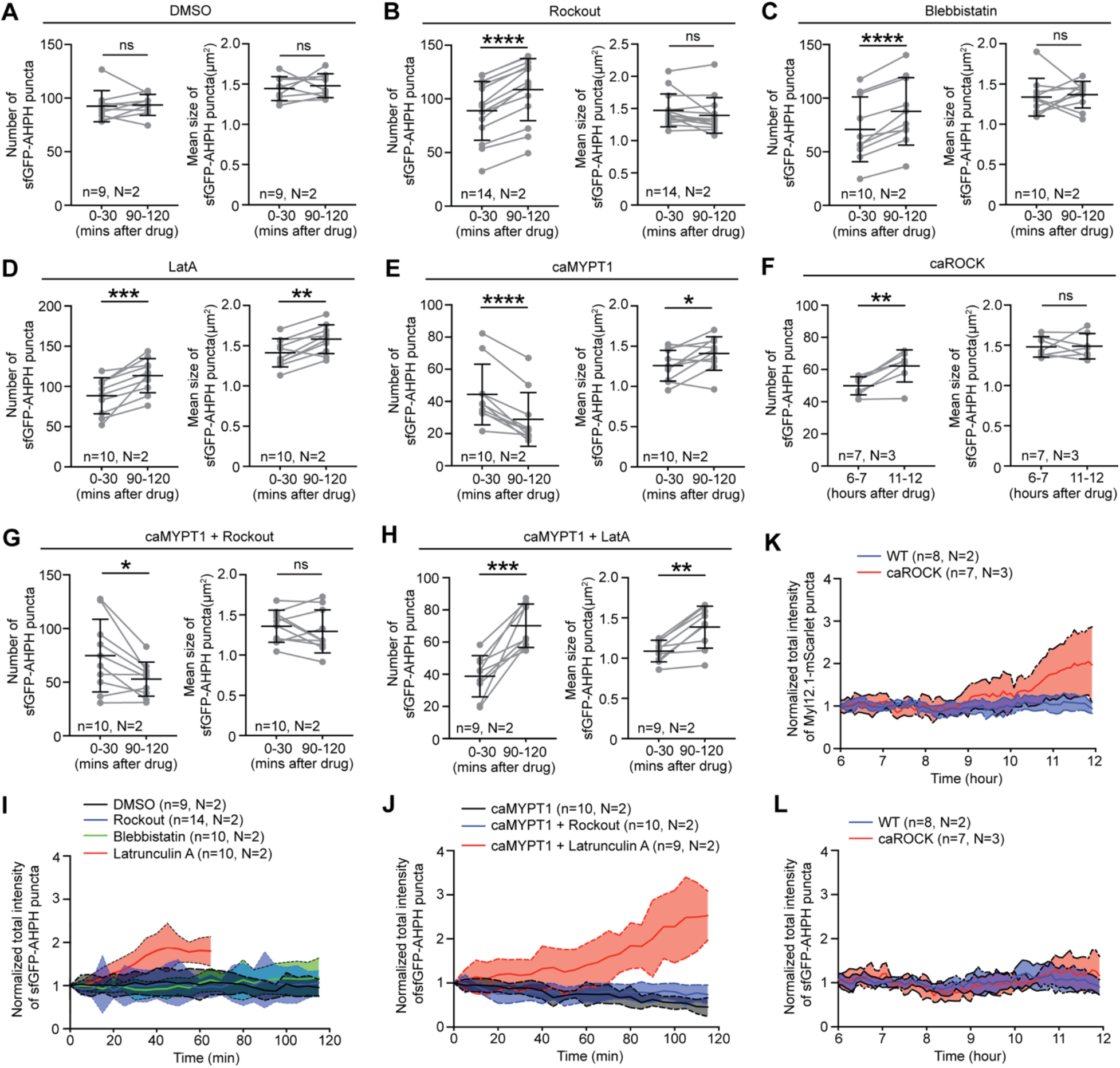
RhoA activation in the primordium of embryos with perturbations in RhoA-mediated signaling. **A–D:** Quantifications of total number and mean size of sfGFP-AHPH puncta in the primordium after treatment with DMSO (A), Rockout (B), blebbistatin (C), or Latrunculin A (D). The time stamp indicates minutes after drug addition. Each dot in the quantification represents the average of 7 time points of the indicated time periods. Images were taken every 5 minutes. Mean and s.d. are indicated, and each pair of grey dots connected by a line represents the same embryo. B-C: \*\*\*\**P* < 0.0001 (Two-tailed paired t test). D: \*\*\**P* = 0.0008 (number of puncta); \*\**P* = 0.0014 (size of puncta) (Two-tailed paired t test). ns = not significant. **E–F**, Quantifications of total number and mean size of sfGFP-AHPH puncta in primordia expressing caMYPT1 (E) or caROCK (F). The time indicates minutes (E) or hours (F) after RU-486 addition to the media. Each dot in the quantification represents the average of 7 (E) or 12 (F) time points of the indicated time periods. Images were taken every 5 minutes. Mean and s.d. are indicated, and each pair of grey dots connected by a line represents the same embryo. E: \**P* = 0.0213 (size of puncta) (Two-tailed paired t test); \*\*\*\**P* < 0.0001 (number of puncta) (Two-tailed paired t test); F: \*\**P* = 0.0025 (number of puncta) (Two-tailed paired t test); ns = not significant. **G–H:** Quantifications of total number and mean size of sfGFP-AHPH puncta in primordia expressing caMYPT1 in embryos treated with Rockout (G) or Latrunculin A (H). The time indicates minutes after drug addition. caMYPT1 expression was induced by adding RU-486 one hour prior to adding the drugs to the embryo media. Each dot in the quantification represents the average of 7 time points of the indicated time periods. Images were taken every 5 minutes. Mean and s.d. are indicated, and each pair of grey dots connected by a line represents the same embryo. G: \**P* = 0.0134 (number of puncta) (Two-tailed paired t test). H: \*\*\**P* = 0.0003 (number of puncta) (Two-tailed paired t test); \*\**P* = 0.0013 (size of puncta) (Two-tailed paired t test). ns = not significant. **I**, Quantifications of the sfGFP-AHPH intensity in the primordium after treatment with DMSO, Rockout, blebbistatin, and Latrunculin A normalized to the 0 min time point. Mean and s.d. are indicated. **J**, Quantifications of the sfGFP-AHPH intensity in the primordium after induction of caMYPT1 in the presence or absence of Rockout or Latrunculin A normalized to the 0 min time point. Mean and s.d. are indicated. **K**, Quantifications of the total Myl12.1-mScarlet intensity in the primordium after induction of caROCK normalized to the 0 min time point. Mean and s.d. are indicated. Note Myl12.1-mScarlet starts to increase in intensity after ∼10 hours of treatment of RU-486. **L**, Quantifications of the total sfGFP-AHPH intensity in the primordium after induction of caROCK normalized to the 0 min time point. Mean and s.d. are indicated. Note that after ∼10 hours of treatment of RU-486, sfGFP-AHPH shows no significant change in intensity. In all panels, n indicates number of primordia/embryos and N indicates number of experiments.

